# Logic-Based Modeling of Inflammatory Macrophage Crosstalk with Glomerular Endothelial Cells in Diabetic Kidney Disease

**DOI:** 10.1101/2023.04.04.535594

**Authors:** Krutika Patidar, Ashlee N. Ford Versypt

## Abstract

Diabetic kidney disease is a complication in one out of three patients with diabetes. Aberrant glucose metabolism in diabetes leads to structural and functional damage in glomerular tissue and a systemic inflammatory immune response. Complex cellular signaling is at the core of metabolic and functional derangement. Unfortunately, the mechanism underlying the role of inflammation in glomerular endothelial cell dysfunction during diabetic kidney disease is not fully understood. Mathematical models in systems biology allow the integration of experimental evidence and cellular signaling networks to understand mechanisms involved in disease progression. This study developed a logic-based ordinary differential equations model to study inflammatory crosstalk between macrophages and glomerular endothelial cells during diabetic kidney disease progression using a protein signaling network stimulated with glucose and lipopolysaccharide. This modeling approach reduced the biological parameters needed to study signaling networks. The model was fitted to and validated against available biochemical data from *in vitro* experiments. The model identified mechanisms for dysregulated signaling in macrophages and glomerular endothelial cells during diabetic kidney disease. In addition, the influence of signaling interactions on glomerular endothelial cell morphology through selective knockdown and downregulation was investigated. Simulation results showed that partial knockdown of VEGF receptor 1, PLC-γ, adherens junction proteins, and calcium partially recovered the intercellular gap width between glomerular endothelial cells. These findings contribute to understanding signaling and molecular perturbations that affect the glomerular endothelial cells in the early stage of diabetic kidney disease.

**NEW & NOTEWORTHY:** The work provides a novel analysis of signaling crosstalk between macrophages and glomerular endothelial cells in the early stage of diabetic kidney disease. A logic-based mathematical modeling approach identified vital signaling molecules and interactions that regulate glucose-mediated inflammation in the glomerular endothelial cells and cause endothelial dysfunction in the diabetic kidney. Simulated interactions among vascular endothelial growth factor receptor 1, nitric oxide, calcium, and junction proteins significantly affect the intercellular gap between glomerular endothelial cells.

## INTRODUCTION

Diabetic kidney disease (DKD) is a major microvascular complication in the kidney that affects about 30% of patients with type 1 diabetes and 40% of patients with type 2 diabetes (1). DKD causes reduced kidney function and may progress into end-stage renal disease (ESRD). Reports suggest that approximately 12–55% of cases of ESRD are due to diabetes (2). This imposes a high burden on the cost of living and quality of life. The high mortality in diabetic patients is often associated with DKD complications and progression to ESRD (3). Although diabetes management is one of the preventative treatments for DKD (4), the rising number of cases of type 1 and type 2 diabetes continues to increase the DKD prevalence (1). There is an urgent need to understand the pathophysiology of DKD better to design effective and preventative treatment strategies that can mitigate early signs of microvascular damage and ultimately reduce DKD prevalence in diabetic patients (3).

Ongoing research suggests that DKD is caused by many complex and interlinked processes associated with high blood glucose and poor glycemic control. The hyperglycemic microenvironment in type 2 diabetes has been linked to metabolic dysfunction in immune cells and inflammation (5–7). The microenvironment influences the activation of pro-inflammatory stimuli, such as exogenous lipopolysaccharide (LPS), interferon-gamma (IFN-γ), and tumor necrosis factor-alpha (TNF-α) (7, 8), which in turn increase glucose uptake in immune cells (9). In the early stage of DKD, immune cells, including macrophages, tend to polarize to a pro-inflammatory phenotype over an anti-inflammatory phenotype (9, 10). The synergistic action of pro-inflammatory mediators and increased glucose uptake contribute to dysregulated immune response and signaling (9). This dysregulated signaling enhances the expression of pro-inflammatory molecules, such as interleukin (IL)-6, TNF-α, IL-1β, and reactive oxygen species (ROS), which affect downstream macrophage response and activity (9, 10). Inflammatory molecules can recruit leukocytes to sites of inflammation and initiate the process of leukocyte transmigration (11, 12). These circulating pro-inflammatory molecules infiltrate the glomerular interstitium (Fig. 1a) in the diabetic kidney (13) and cause microvascular leakage by upregulation and redistribution of cell-cell adhesion molecules (8, 11, 14, 15). Recruitment of inflammatory macrophages and their mediators is a critical event in the pathophysiology of the early stage of DKD (13). There is still an incomplete understanding of the highly complex and diverse molecular interactions in the diabetic kidney (8). Hence, a deeper analysis of glucose-mediated inflammatory mechanisms and pathways in the diabetic kidney can contribute to a more comprehensive understanding of DKD development and progression.

**Figure 1:**
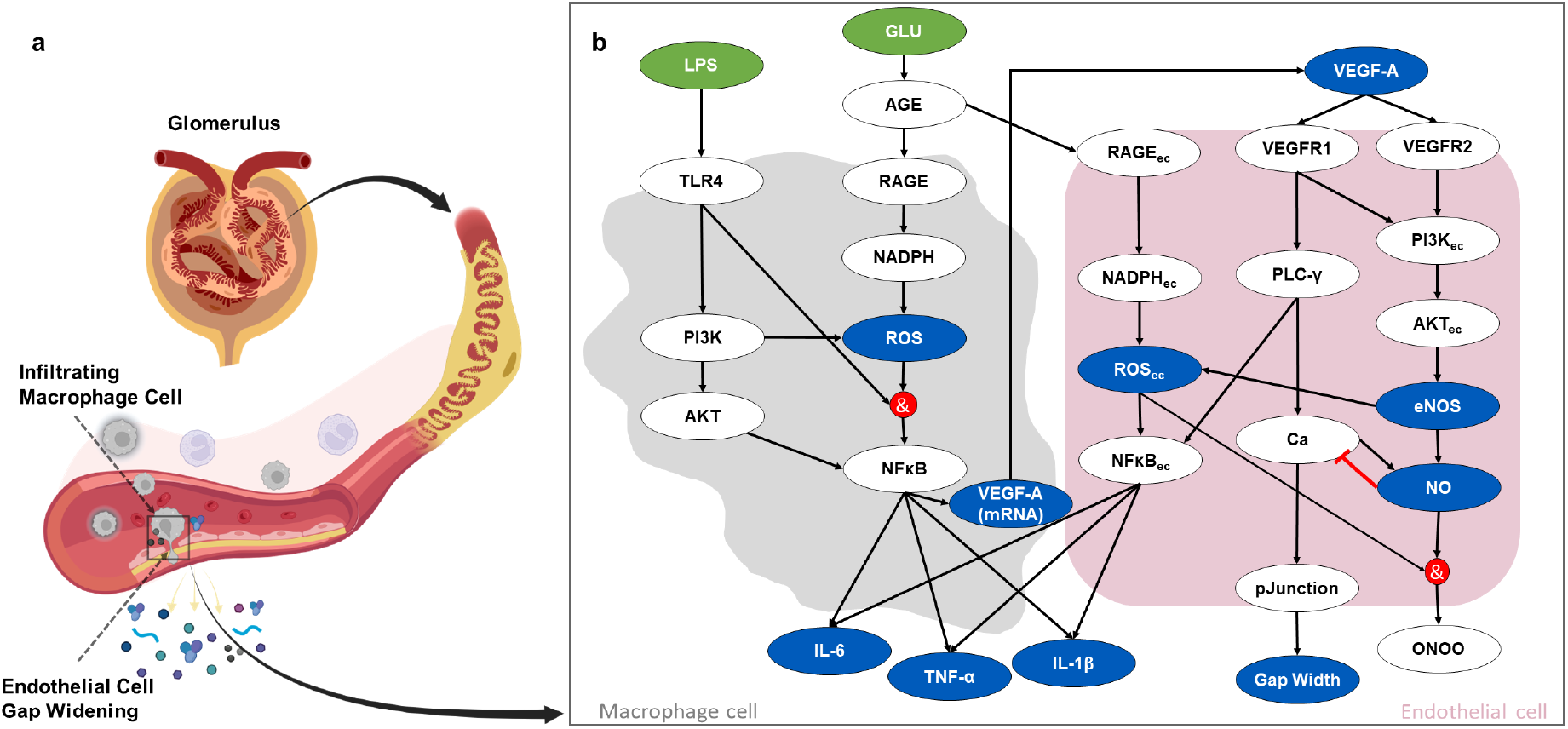
(a) Visualization of immune-mediated signaling dysregulation and dysfunction in the glomerulus. Infiltrating macrophage cells interact with glomerular endothelial cells, which experience a widening of the intercellular gaps between endothelial cells, leading to the loss of solute and macromolecules from the blood. Created with BioRender.com. (b) The multi-cellular protein signaling network of crosstalk between macrophages (left) and glomerular endothelial cells (right) is stimulated with glucose (GLU) and lipopolysaccharide (LPS), a pro-inflammatory stimulus. Green ovals are input nodes, blue ovals are output nodes, and white ovals are regulatory nodes. Black arrows are activating interactions, a red line with a flat-head arrow is an inhibiting interaction, and red circles indicate logic *AN D* gates. An *OR* logic rule connects two or more edges to a subsequent node throughout the network unless indicated otherwise by an *AN D* logic gate. The subscript “ec” denotes an intracellular species expressed in endothelial cells. IL-6, TNF-α, IL-1β, and VEGF-A are protein levels expressed in extracellular space. ROS, ROS_ec_, VEGF-A_mRNA_, and NO are expressed within the cells. The Gap Width node denotes a fractional change in the intercellular gap between GECs. The pJunction node represents the phosphorylated junction protein levels. Abbreviations are defined in Table 1.

Glomerular endothelial dysfunction is a pathological phenomenon that characterizes the early stages of DKD development and progression. Glomerular endothelial cells (GECs) are early targets of diabetes-dependent inflammation and injury (2). Glomerular endothelial activation is accompanied by changes in the expression of leukocytes, cytokine production, and loss of barrier integrity (16). Numerous studies suggest that activation of GECs in early DKD can dysregulate signaling pathways, promote intracellular hyperglycemia, elevate advanced glycation end products (AGEs) and their receptors, affect cell-cell adhesion, increase glomerular permeability, and promote the recruitment of inflammatory molecules (17). LPS-stimulated pro-inflammatory macrophages are involved in the activation of GECs *in vitro* (18). The inflammatory mediators and immune cells disrupt vascular endothelial growth factor (VEGF) expression, which plays a central role in maintaining the vascular function of GECs and nitric oxide (NO) homeostasis (19, 20). Imbalanced VEGF expression causes uncoupling of endothelial nitric oxide synthase (eNOS), imbalance of NO (19), and loss of endothelial barrier integrity (11, 12). The uncoupling of eNOS causes eNOS to produce NO as a product and ROS as a by-product, which leads to excessive oxidation of NO and reduces NO bioavailability in GECs (21). Hence, the uncoupling of eNOS is a crucial indicator of GEC activation and dysfunction. NO bioavailability within GECs has been associated with glomerular endothelial permeability (22, 23). An association between NO bioavailability and endothelial cell junction integrity has been suggested (11, 24). In the kidneys, hyperglycemia and inflammatory stimuli induce intercellular junction disruption and enhanced microvascular leakage in the glomeruli. The loss of intercellular junction integrity leads to increased paracellular permeability, leukocyte transmigration, and dysregulated intracellular signals (11, 15). Many experimental studies have associated VEGF-mediated tyrosine phosphorylation of junction proteins with increased permeability (25, 26), decreased transendothelial electrical resistance (27, 28), and regulating the passage of solutes/macromolecules in kidney (27–29). Stimulation with the inflammatory cytokine IL-6 affected junction protein VE-cadherin, and phosphorylation increased solute permeability and reduced transendothelial electrical resistance in GECs (12). Thus, the intercellular gap width changes are considered an early marker of GEC activation and dysfunction during DKD progression (Fig. 1a). About 50% of the filtration through the glomerulus is dependent on GECs and the endothelial cell surface layer, making GECs an essential contributor to the glomerular permeability of the kidney (30). Transcellular and paracellular routes may restrict the transport of water, solutes, or macromolecules through GECs. The fenestrations are transcellular holes located throughout GECs and are among the main contributors to the glomerular filtration barrier (31, 32). Despite some experimental studies on GECs in healthy and diseased states, a direct mechanistic link between macrophage-dependent GEC activation and dysfunction in the early stage of DKD is not fully understood (24). This study investigated the early pathological events under hyperglycemic and inflammatory conditions leading to GEC activation and signaling dysregulation.

**Table 1:** Chemical species abbreviations and definitions used in the study.

| Abbreviation | Definition |
| --- | --- |
| AGE | advanced glycation end product |
| AKT | serine/threonine-specific protein kinases |
| Ca | calcium |
| eNOS | endothelial nitric oxide synthase |
| Gap Width | intercellular gap width between GECs |
| GLU | glucose |
| IL | interleukin |
| LPS | lipopolysaccharide |
| NADPH | nicotinamide adenine dinucleotide phosphate |
| NFκB | nuclear factor kappa B |
| NO | nitric oxide |
| ONOO | peroxynitrite |
| PI3K | phosphoinositide 3-kinases |
| pJunction | phosphorylated junction protein |
| PLC-γ | phospholipase C gamma |
| RAGE | receptor of advanced glycation end product |
| ROS | reactive oxygen species |
| TLR | toll-like receptor |
| TNF-α | tumor necrosis factor-alpha |
| VEGF | vascular endothelial growth factor |
| VEGFR | vascular endothelial growth factor receptor |

In the present study, we built a protein signaling network (PSN) of interconnected pathways between two cell types—GECs and macrophages—to study glucose-mediated inflammatory mechanisms in DKD. A PSN depicts the relationship between interacting proteins using directional arrows (33). The PSN used here was assembled manually using published literature for selected species and pathways in the PSN and curated experimental data; we have also explored using artificial intelligence-enabled text mining for automated PSN assembly in other work (34). Here, we used a mathematical approach to integrate published experimental evidence and the PSN. Ordinary differential equations (ODEs) are often used to model PSNs and other biochemical networks (35–38), but ODE-based models rely on extensive data, parameter values, and information on reaction kinetics. Alternatively, modelers have used logic-based modeling, fuzzy logic modeling, and extreme pathways analysis to reduce the burden of kinetics information needed to study complex mammalian signaling networks (33, 39). These techniques require few or no parameters. For instance, a discrete logic model (Boolean model) successfully showed the changes in endothelial cell behavior and signaling due to extracellular microenvironments (40). A logic-based model (LBM) was used to gain a qualitative understanding of species expression, signaling pathways, and missing interactions involved in T cell activation (41). Furthermore, pathway analysis of differentially expressed genes and profiles in early DKD has pointed to critical pathophysiological factors in DKD (42). However, each approach has limitations, including a lack of prediction capability for continuous dynamics (time course) in Boolean models, unrealistic timescales in fuzzy logic analysis, and extensive data collection requirements for pathway analysis.

Here, the emphasis was on predicting continuous signaling dynamics while incorporating the prior mechanistic knowledge of intracellular reaction details and the limited available kinetic data. Thus, we considered mathematical methods that combine ODEs and LBMs to leverage the advantages of each approach. Previously, a hybrid modeling method used an ODE model to explain pathway interactions and a qualitative logic-based model to define cell-cell interactions within a tissue environment (43). We adopted a modeling technique that combines continuous dynamic ODE models with qualitative LBMs (39), termed a logic-based ordinary differential equations (LBODEs) model. Several research studies incorporated such transformation of logic-based models into systems of first-order ODEs (44, 45). One study transformed a discrete Boolean model into ODEs using non-normalized Hill functions in the Odefy toolbox to study ligand concentrations and binding affinities for the activation of T cells (44). Logic-based formalism was also used to reverse engineer complex biochemical networks from data and prior knowledge using metaheuristic optimization strategies, as seen in CellNOpt software (46, 47). Netflux is another user-friendly software for developing dynamic logic-based mathematical models of biological networks (39). Kraeutler et al. presented the advantages of using LBODEs, mainly when biochemical parameters are limited or unavailable; LBODEs were shown to identify unexpected network relationships, to generate experimentally testable predictions, and to agree with findings from a validated biochemical model (39). The Netflux software is readily available and easy to use for building a PSN and generating the corresponding set of LBODEs that comprise the mathematical model for a biochemical system. Such tested software standardizes the methods used to obtain biological insights, which may not be evident through discrete models (44). We used Netflux to apply the LBODEs modeling approach to our PSN between macrophages and GECs stimulated under different conditions to analyze the effects of the signaling crosstalk between these cell types. We predicted species dynamics, network relationships, and critical drivers of macrophage-dependent inflammation and glomerular endothelial dysfunction in the early stage of DKD.

## METHODS

### Protein signaling network assembly

We formulated a PSN for crosstalk between GECs and macrophages (Fig. 1b) by manually assembling information from literature studies (8, 9, 19, 21, 40, 48–54) and pathway databases (55, 56). The PSN captures the signaling pathways and molecules (Table 1) that cause inflammation and dysfunction in two cell types: macrophages and GECs. The multi-cellular PSN has 30 species nodes and 40 interaction edges for reactions (Fig. 1b). The green ovals are input nodes or stimuli, the white ovals are regulatory nodes, and the blue ovals are model response output nodes. The network has two stimuli: glucose (GLU) and LPS. LPS is used as a representative systemic inflammatory stimulus like that triggered by diabetes. The *in vitro* data for model fitting used LPS as an exogenous inflammatory stimulus as it is known to induce a systemic immune response in humans (57). The network is stimulated with three different treatment conditions: GLU only, LPS only, and both GLU and LPS. Based on the logic-based formalism, two or more incoming arrows to a node are connected by an *OR* logic rule except where a red circle is placed denoting an *AN D* logic rule (Fig. 1b). An *OR* logic rule between two input nodes is used when either input can activate (black arrow) or inhibit (red flat-head arrow) the output node. The *OR* logic combines any number of input nodes acting on the same output; the corresponding reaction rules are listed independently but are joined by the *OR* operator in the LBODEs. An *AN D* logic rule between two input nodes is used when both inputs are needed to activate the output node; a single reaction rule is defined for an *AN D* logic rule. Fig. 2 shows representative examples of the interaction types between nodes in the PSN from Fig. 1b. Considering the complex nature of network interactions, we included signaling pathways involved in two main events in early DKD. These events include the release of pro-inflammatory molecules due to the infiltration of inflammatory macrophages and macrophage-dependent GEC activation and dysfunction. The rationale behind selecting these pathways and molecules is discussed further.

**Figure 2:**
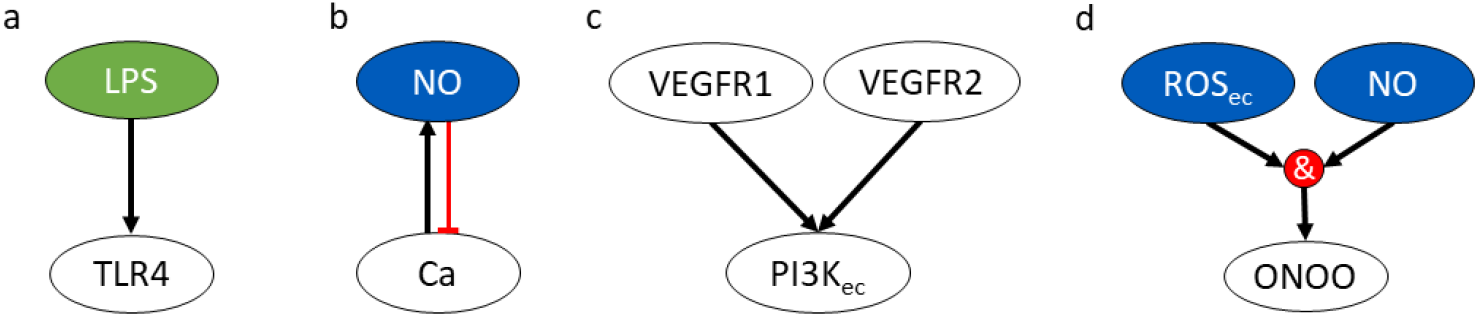
Logic-based formalism for a protein signaling network (PSN). The species interactions (a)–(d) are used as examples to explain logic-based reaction rules for the interaction edges in the PSN (Fig. 1). Each of the example descriptions is followed by its corresponding reaction rule. (a) LPS activates TLR4: LPS ⇒TLR4. (b) NO inhibits Ca: !NO⇒ Ca, and Ca activates NO: Ca ⇒NO. (c) VEGFR1 or VEGFR2 activate PI3K_ec_ using an *OR* logic rule: VEGFR1 ⇒PI3K_ec_ *OR* VEGFR2⇒ PI3K_ec_. (d) ROS_ec_ and NO activate ONOO using an *AN D* logic rule: ROS_ec_ & NO ⇒ONOO. A black arrow is an activating interaction, a red line with a flat-head arrow is an inhibiting interaction, and a red circle is a logic *AN D* gate. The subscript “ec” denotes an intracellular species expressed in endothelial cells. Abbreviations are defined in Table 1.

In diabetes, immune cells like macrophages fail to metabolize glucose or regulate the efficient transport of glucose, which leads to intracellular hyperglycemia (58, 59). Studies showed that the hyperglycemic microenvironment initiates a pro-inflammatory signaling response downstream, mediated by the activation of LPS and toll-like receptors (TLR) such as TLR4 (9, 10). In cell culture experiments, bacterial LPS is a representative pathogen-associated molecular pattern (PAMP) that allows mammalian cells to recognize bacterial invasion and trigger innate immune responses (9). In cultured macrophages, LPS stimulates a robust inflammatory response, which initiates glucose-mediated inflammation in macrophages (9). Hence, LPS is considered one of the stimuli in our PSN.

LPS transduction signal is carried by activation of PI3K and AKT kinases, which mediate macrophage response, energy consumption, and energy metabolism (60). AKT is a primary substrate of PI3K and is associated with regulating inflammatory and energetic metabolic responses. Therefore, we speculate that PI3K and AKT are crucial in macrophage polarization in the PSN (9).

Advanced glycation is one of the major pathways involved in the development and progression of different diabetic complications, including DKD (54). This protein glycation leads to advanced glycation end products (AGE), which activate and modify intracellular pathways through receptor-mediated signaling (54). The expression of AGE receptors (RAGE) is enhanced in cells like macrophages and endothelial cells during diabetes and inflammation. AGE and RAGE interaction causes oxidative stress and activation of nuclear factor kappa B (NFκB) (54). The glycated proteins release ROS via activation of NADPH oxidase (49, 61). Moreover, aberrant PI3K-AKT signaling increases ROS levels through either direct modulation of mitochondrial bioenergetics or indirect production of ROS as a metabolic by-product (62). Hence, NADPH- and PI3K-dependent ROS are considered in the PSN to gain a better perspective on mediators of ROS.

Further, transcription factor NFκB contributes to the structural and functional changes observed in DKD, including pro-inflammatory macrophage polarization (54). NFκB is responsible for the expression of pro-inflammatory cytokines, like IL-6, TNF-α, and IL-1β, and other inflammatory markers, growth factors including VEGF-A, and adhesion molecules (8).

The macrophage-dependent VEGF-A expression is associated with the direct activation of GECs in the PSN. VEGF expression modulates the crosstalk between endothelial cells and macrophages (63). VEGF and its isoforms are critical to GECs. Excess VEGF contributes to DKD development and progression through disrupted signaling in GECs, whereas low VEGF levels can result in loss of capillaries and endothelial cells in the glomerulus (64). VEGF expression promotes endothelial cell development through NO modulation (64). Studies suggested that glucose indirectly regulates VEGF-A and the signaling of its receptors (VEGFR1 and VEGFR2) in GECs (63, 65, 66). VEGFR1 is a positive regulator of inflammation and immune cell recruitment via PLC-γ and PI3K pathways (51, 67, 68). The downstream signaling of VEGFR1 has yet to be fully understood, mainly due to its mild biological activity in culture (51). VEGF-A mediates signal transduction for cell proliferation, survival, migration, and enhanced vascular permeability via VEGFR2 (20). VEGFR2-dependent signaling promotes eNOS activity, which produces NO, an important vasodilator in GECs (20). VEGF-A and VEGFR2 activity reduced in the GECs during diabetes (69). This phenomenon leads to perturbations in downstream signaling, eNOS uncoupling, and reduced NO bioavailability (70). As a result, excess ROS in GECs can react with available NO to form reactive nitrogen species like peroxynitrite (ONOO) (64, 71). In the PSN, the VEGF-dependent signaling pathway accounts for VEGF-A receptors VEGFR1 and VEGFR2, PLC-γ, eNOS, NO, ROS, and reactive nitrogen species expression. The VEGF-dependent NO bioavailability is connected with GEC dysfunction.

Cell-cell junction proteins hold adjacent GECs together and modulate the passage of macromolecules, proteins, or infiltrating macrophages through the GEC layer. For activated GECs, the phosphorylation of these junction proteins, such as VE-cadherin, is regulated by calcium (Ca) expression via the PLC-γ pathway and NO expression (23, 72). The phosphorylation of junction proteins like VE-cadherin causes a loss of barrier integrity in GECs and triggers the opening of the intercellular junctions. Intercellular junction protein is shown as a pJunction node in the PSN. We posit that the junction opening or closing is directly proportional to the size of the intercellular gap between GECs, shown as the Gap Width node in the network.

In addition to the literature-based prior knowledge discussed above that is incorporated into the PSN, the following list summarizes additional assumptions and restrictions in scope for the PSN:

- Literature suggests a higher polarization of the pro-inflammatory phenotype of macrophage cells during early stages of DKD (9). We considered a pro-inflammatory phenotype of macrophages expressing pro-inflammatory species and ignored anti-inflammatory species for simplicity.
- GECs comprise a negatively charged matrix, the glycocalyx, which also contributes to endothelial permeability (2). We only focused on the endothelial cell layer for GEC signaling and dysfunction. Glycocalyx signaling and characteristics were not part of this study.
- AGEs are mediators and drivers of diabetic complications (18, 73). We assumed that glucose directly regulates AGE activation in the PSN. The mechanism regulating glucose-dependent AGE activation is not part of this study.
- The interaction kinetics across different endothelial cell lines are assumed to be the same, but protein levels of species may differ across cell lines (20). This has been previously considered in other models where data from different endothelial cell lines were curated due to the lack of studies on a specific cell line.
- The protein signaling network considers only two types of cells: macrophages and GECs. These cells are stimulated by GLU and LPS, which mimic hyperglycemic and inflammatory responses during the early stages of DKD, respectively.

Using a fixed physiological range allowed us to extract protein expression data from studies performed under similar treatment conditions. We assumed that full activation of GLU stimulus (i.e., *W*_GLU_ = 1) is analogous to a 25–30 mM hyperglycemic range. Full activation of LPS (*W*_LPS_ = 1) is the same as an LPS treatment of 0.1 µg/ml in cultured cells.

### Logic-based ordinary differential equations model

We simulated the dynamic influences of stimuli through the PSN using an LBODEs model. The LBODEs modeling technique combines ODEs that are continuous functions of time with qualitative logic-based Boolean up-or down-regulation (i.e., activation or inhibition) using normalized Hill functions (saturating sigmoidal terms) for the logic-based modeling portion (39). The LBODEs model equations were generated by the Netflux software package (74) after we supplied the necessary interaction rules and species list from the PSN that we assembled (Tables 2 and 3). The Netflux package was used to generate the LBODEs model equations and default parameters, which were exported to MATLAB (75). MATLAB was used to simulate, fit, and validate the model with further customization beyond Netflux for parameter estimation and other analyses. The LBODEs were solved using the ode23s function in MATLAB.

**Table 2:** List of optimal reaction parameters for the LBODEs model organized by reaction index *j* . Reaction weight (*W*_*j*_), Hill coefficient (*n*_*j*_), and half-maximal 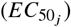 are shown for respective reaction rules or interactions. Reaction rule A⇒ C denotes input A activates output C. Reaction rule !A⇒ C denotes input A inhibits output C. Reaction rule A & B C denotes the *AN D* logic operator between input A and input B to give output C. Default values are *W*_*j*_ = 1, *n*_*j*_ = 1.4, and 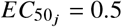.Estimated values of sensitive parameters are displayed with three significant figures. *W*_*j*_ = 0 for ⇒GLU and ⇒LPS reaction rules represent inactive states. *W*_*j*_ is set to 1 based on the active treatment condition for *j* = 1 and/or *j* = 2. *W*_1_ and *W*_2_ are referred as *W*_GLU_ and *W*_LPS_, respectively. The source terms for GLU (*s* (GLU)) and LPS (*s* (LPS)) are only dependent on *W*_*j*_ and independent of *n*_*j*_ and 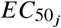.Abbreviations are defined in Table 1.

| Index $j$ | Reaction Rule | $W_j$ | $n_j$ | $EC_{50_j}$ |
| --- | --- | --- | --- | --- |
| 1 | ⇒ GLU | 0 | - | - |
| 2 | ⇒ LPS | 0 | - | - |
| 3 | LPS ⇒ TLR4 | 1 | 2.70 | 0.419 |
| 4 | GLU ⇒ AGE | 1 | 1.44 | 0.5 |
| 5 | AGE ⇒ RAGE | 1 | 2.71 | 0.474 |
| 6 | RAGE ⇒ NADPH | 0.944 | 2.69 | 0.470 |
| 7 | TLR4 & ROS ⇒ NFκB | 1 | 2.69 | 0.5 |
| 8 | TLR4 ⇒ PI3K | 0.950 | 2.69 | 0.422 |
| 9 | NADPH ⇒ ROS | 0.943 | 2.63 | 0.545 |
| 10 | PI3K ⇒ AKT | 1 | 2.69 | 0.5 |
| 11 | PI3K ⇒ ROS | 0.950 | 2.70 | 0.419 |
| 12 | NFκB<sub>ec</sub> ⇒ TNF-α | 0.999 | 3.96 | 0.839 |
| 13 | AKT ⇒ NFκB | 1 | 2.70 | 0.5 |
| 14 | NFκB ⇒ IL-6 | 0.950 | 2.70 | 0.420 |
| 15 | NFκB ⇒ TNF-α | 0.950 | 2.70 | 0.419 |
| 16 | NFκB ⇒ VEGF-A<sub>mRNA</sub> | 0.949 | 2.70 | 0.422 |
| 17 | VEGF-A<sub>mRNA</sub> ⇒ VEGF-A | 0.948 | 2.77 | 0.594 |
| 18 | NFκB ⇒ IL-1β | 0.950 | 2.69 | 0.421 |
| 19 | VEGF-A ⇒ VEGFR1 | 1 | 2.71 | 0.5 |
| 20 | VEGF-A ⇒ VEGFR2 | 1 | 2.72 | 0.5 |
| 21 | AGE ⇒ RAGE<sub>ec</sub> | 1 | 1.55 | 0.838 |
| 22 | RAGE<sub>ec</sub> ⇒ NADPH<sub>ec</sub> | 0.937 | 1.59 | 0.838 |
| 23 | VEGFR2 ⇒ PI3K<sub>ec</sub> | 1 | 2.72 | 0.5 |
| 24 | VEGFR1 ⇒ PI3K<sub>ec</sub> | 1 | 2.72 | 0.5 |
| 25 | NADPH<sub>ec</sub> ⇒ ROS<sub>ec</sub> | 0.946 | 3.91 | 0.838 |
| 26 | PI3K<sub>ec</sub> ⇒ AKT<sub>ec</sub> | 1 | 2.92 | 0.688 |
| 27 | AKT<sub>ec</sub> ⇒ eNOS | 1 | 3.12 | 0.576 |
| 28 | VEGFR1 ⇒ PLC-γ | 1 | 1.4 | 0.5 |
| 29 | PLC-γ ⇒ NFκB<sub>ec</sub> | 1 | 1.4 | 0.5 |
| 30 | ROS<sub>ec</sub> ⇒ NFκB<sub>ec</sub> | 1 | 2.70 | 0.0100 |
| 31 | NFκB<sub>ec</sub> ⇒ IL-6 | 0.986 | 2.69 | 0.471 |
| 32 | NFκB<sub>ec</sub> ⇒ IL-1β | 0.962 | 2.70 | 0.391 |
| 33 | eNOS ⇒ NO | 1 | 1.45 | 0.156 |
| 34 | eNOS ⇒ ROS<sub>ec</sub> | 1 | 3.66 | 0.832 |
| 35 | ROS<sub>ec</sub> & NO ⇒ ONOO | 1 | 1.4 | 0.5 |
| 36 | !NO ⇒ Ca | 1 | 2.68 | 0.5 |
| 37 | PLC-γ ⇒ Ca | 1 | 1.4 | 0.5 |
| 38 | Ca ⇒ pJunction | 1 | 1.4 | 0.5 |
| 39 | pJunction ⇒ Gap Width | 1 | 1.4 | 0.5 |
| 40 | Ca ⇒ NO | 1 | 1.4 | 0.5 |

**Table 3:** List of optimal species parameter τ_*i*_ for the LBODEs model organized by species index *i*. The time constant (τ_*i*_) values in units of hours for the species used in the model are shown for each treatment condition. The default value of τ_*i*_ is 1 hour. Estimated values of sensitive parameters are displayed with three significant figures. The default values of other species parameters are the initial value 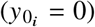 and maximal value 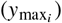 for all species unless stated otherwise. 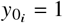 for eNOS, NO, VEGF-A_mRNA_, and VEGF-A species for the GLU only treatment condition. Abbreviations are defined in Table 1.

| Index $i$ | Species | GLU only | LPS only | GLU and LPS |
| --- | --- | --- | --- | --- |
| 1 | GLU | 1 | 1 | 1 |
| 2 | LPS | 5.05 | 0.800 | 1.00 |
| 3 | AGE | 1 | 1 | 1 |
| 4 | VEGFR1 | 1 | 1 | 1 |
| 5 | VEGFR2 | 1 | 1 | 1 |
| 6 | VEGF-A<sub>mRNA</sub> | 3.38 | 0.944 | 1.47 |
| 7 | RAGE<sub>ec</sub> | 1 | 1 | 1 |
| 8 | RAGE | 1 | 1 | 1 |
| 9 | TLR4 | 5.03 | 0.800 | 1.00 |
| 10 | NADPH | 1 | 1 | 1 |
| 11 | NADPH<sub>ec</sub> | 1 | 1 | 1 |
| 12 | ROS<sub>ec</sub> | 9.96 | 1.06 | 5.15 |
| 13 | ROS | 5.88 | 0.800 | 1.00 |
| 14 | PI3K | 5.03 | 0.800 | 1.00 |
| 15 | AKT | 1 | 1 | 1 |
| 16 | PI3K<sub>ec</sub> | 6.59 | 0.977 | 1.00 |
| 17 | AKT<sub>ec</sub> | 5.75 | 0.984 | 1.00 |
| 18 | NFκB<sub>ec</sub> | 1 | 1 | 1 |
| 19 | NFκB | 5.06 | 0.800 | 1.00 |
| 20 | NO | 4.35 | 1.00 | 6.80 |
| 21 | ONOO | 1 | 1 | 1 |
| 22 | eNOS | 1.91 | 0.981 | 1.00 |
| 23 | IL-6 | 4.65 | 0.800 | 1.00 |
| 24 | TNF-α | 9.89 | 0.800 | 1.00 |
| 25 | IL-1β | 4.90 | 1.00 | 4.94 |
| 26 | PLC-γ | 1 | 1 | 1 |
| 27 | VEGF-A | 3.14 | 0.971 | 1.10 |
| 28 | pJunction | 1 | 1 | 1 |
| 29 | Ca | 1 | 1 | 1 |
| 30 | Gap Width | 1 | 1 | 1 |

Each LBODE defining the rate of change of normalized activity of a species A with index *i* follows the same general form (39):

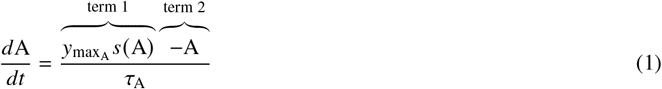

where term 1 is the net activation rate (detailed further below), term 2 is the first-order de-activation rate, and τ_A_ is the characteristic time constant for the species. The initial condition is 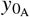 The net activation rate is scaled by 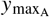,the maximum normalized activation of species A. Note that when discussing the species parameters more generally, we use the notation *τ_i_*, 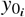, and 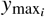.

The quantity *s* (A) is the source of A from activating or inhibiting interactions that act alone or in combination through *OR* or *AN D* logic operators. The activation or inhibition of output node species A is based on an interaction rule computed using a normalized-Hill function dependent on the signal from input node X. The activation function 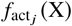 for how species X activates reaction *j* that produces A is

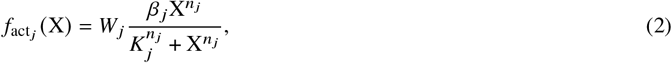

and the inhibition function 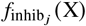 for how species X inhibits reaction *j* that produces A is

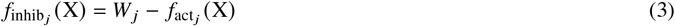

where *W*_*j*_ is the weight of reactionn *j, n*_*j*_ is the Hill coefficient for the reaction, 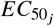 is the half effect,

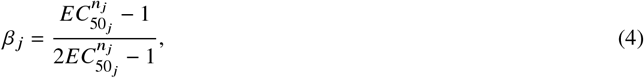

And

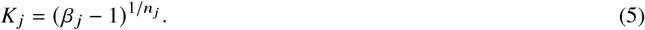

Multiple input signals to the same output node are automatically combined using an *OR* logical operator unless indicated explicitly by an *AN D* logical operator. Mathematically, *OR*(*a, b*) is an *OR* logical operator for input *a* and *b*:

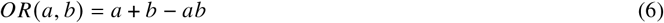

*AN D*(*a, b*) is an *AN D* logical operator for input *a* and *b*:

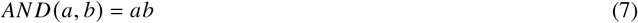

We next illustrate the development of a representative subset of the equations (*Eqs. 8 – 11*) for the nodes and interactions illustrated in Fig. 2. The complete set of interaction rules is provided in Table 2, and the corresponding equations are listed in *Eqs. A1 –A30*.

is A representative single activation interaction rule is LPS ⇒ TLR4 (Fig. 2a). The corresponding LBODE for tracking TLR4 is

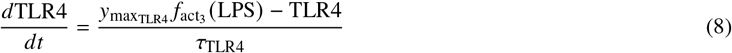

where the source *s* TLR4 is the activation of TLR4 by LPS in reaction rule *j* = 3 (Table 2), which follows *Eq. 2*.

A representative single inhibition interaction rule is !NO ⇒ Ca (Fig. 2b), part of the negative feedback loop between NO and Ca. The corresponding LBODE for tracking Ca is

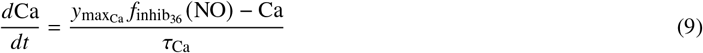

where the source *s* (Ca) is the inhibition of Ca by NO in reaction rule *j* = 36 (Table 2), which follows *Eq. 3*.

If a species is activated by two or more species, an *AN D* or an *OR* logic operator is applied to combine the signals. A representative *OR* logic rule combining two activating signals is VEGFR1 ⇒ PI3K_ec_ *OR* VEGFR2 ⇒ PI3K_ec_ (Fig. 2c). The corresponding LBODE for tracking PI3K_ec_ is

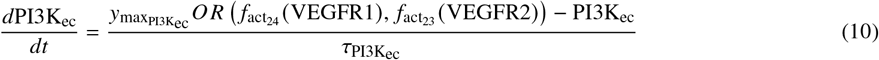

where the source *s* PI3K_ec_ is the *OR* function (*Eq. 6*) combining activation of reaction rules *j* = 23 and *j* = 24 (Table 2), which follow *Eq. 2* individually.

A representative *AN D* logic rule combining two activating signals is ROS_ec_ & NO ⇒ ONOO (Fig. 2d). The corresponding LBODE for tracking ONOO is

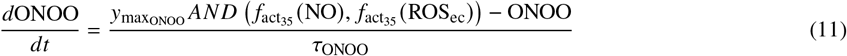

where the source *s* ONOO is the *AN D* function (*Eq. 7*) combining activation by both ROS_ec_ and NO in reaction rule *j* = 35 (Table 2), which follow *Eq. 2* individually.

### Model parameter defaults and constraints

The LBODEs model has parameters that can be divided into two categories: reaction parameters (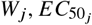 and *n*_*j*_) and species parameters (τ_*i*_, 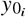, and 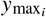). The default values for these parameters, as generated by Netflux, are 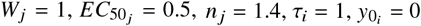 and 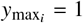.

Reaction parameters include reaction weight *W*_*j*_, half maximal effective concentration or half effect 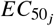,and Hill coefficient *n*_*j*_ . The reaction parameters govern the interactions between species in the network. The parameter definitions, default values, and constraint ranges are used as defined previously (39, 76). For a given interaction, the reaction weight *W*_*j*_ denotes the fraction of activation or inhibition of output species, and 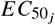 is the fraction of input required to induce a half-maximal effect on the output (39). 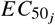 value below 0.5 indicates that less than 50% of input species is required to achieve maximum output activity. Correspondingly, 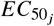 value above 0.5 indicates that more than 50% of input species is required to achieve maximum output activity. The Hill coefficient *n*_*j*_ is a shape factor in the LBODEs model (77). These reaction parameters define the normalized Hill activation and inhibition functions (*Eqs. 2 – 3*). These functions are constrained as *f* (0) = 0, *f* (1) = *W*_*j*_, and *f* (0.5) = 0.5*W*_*j*_ in Netflux (39, 74). Each parameter’s lower and upper bounds are based on prior knowledge, as in Cao et al. (76). The reaction weight constraints are default: 0≤ *W*_*j*_ ≤1. The constraints on 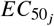 and *n*_*j*_ are selected to satisfy *β* _*j*_ > 1 (*Eqs. 4 – 5*) (76), giving 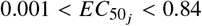 and 1.4 ≤*n*_*j*_ ≤4.

The species parameters include time constant τ_*i*_ (constrained 0.01≤ τ_*i*_ ≤10), initial value 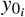, and maximal fractional activation of species *y*_max_ to describe the activity of a species in the network. The time constant (τ_*i*_) parameter is an arbitrary parameter that modulates the signal propagation during early or late events and provides a rough estimate of the local dynamic behavior of the network (41). 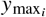 is set to 1 by default for the full activity of a species. 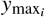 can be below or above a value of 1 to denote species knock-down or over-expression, respectively (39). 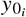 is set to 0 for all species except when species downregulation is predicted. In those cases, 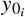 is set to 1.

*W*_*j*_ = 0 for a reaction *j* denotes an inactive state. The source terms for the LBODEs model inputs GLU (*s*(GLU)) and LPS (*s*(LPS)) are only dependent on reaction weight *W*_*j*_ and independent of *n*_*j*_ and 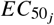.*W*_1_ and *W*_2_ are referred as *W*_GLU_ and *W*_LPS_, respectively, hereafter. Values of *W*_GLU_ or *W*_LPS_ = 1 mean the corresponding input is active.

### Data selection and normalization

The data for model fitting (parameter estimation) and validation were curated from published experimental studies based on the following inclusion criteria:

- co-culture *in vitro* studies (18) were used to study cell-cell interaction between macrophages and GECs with some exceptions (9, 10, 29, 78–84) for individual cell culture studies on either macrophages or endothelial cells,
- bone-marrow derived macrophages (BMDM) from diabetic subjects and the RAW 264.7 murine macrophage cell line were most commonly considered (9, 10, 18, 78, 85),
- cells were stimulated with GLU concentration 5–30 mM and LPS concentration 0–0.1 µg/ml,
- protein concentration from bar graph or data tables at several time points (3, 6, 12, 24, and 48 hours) for three treatment conditions GLU only, LPS only, and both GLU and LPS were collected where available (e.g., Fig. 3).

**Figure 3:**
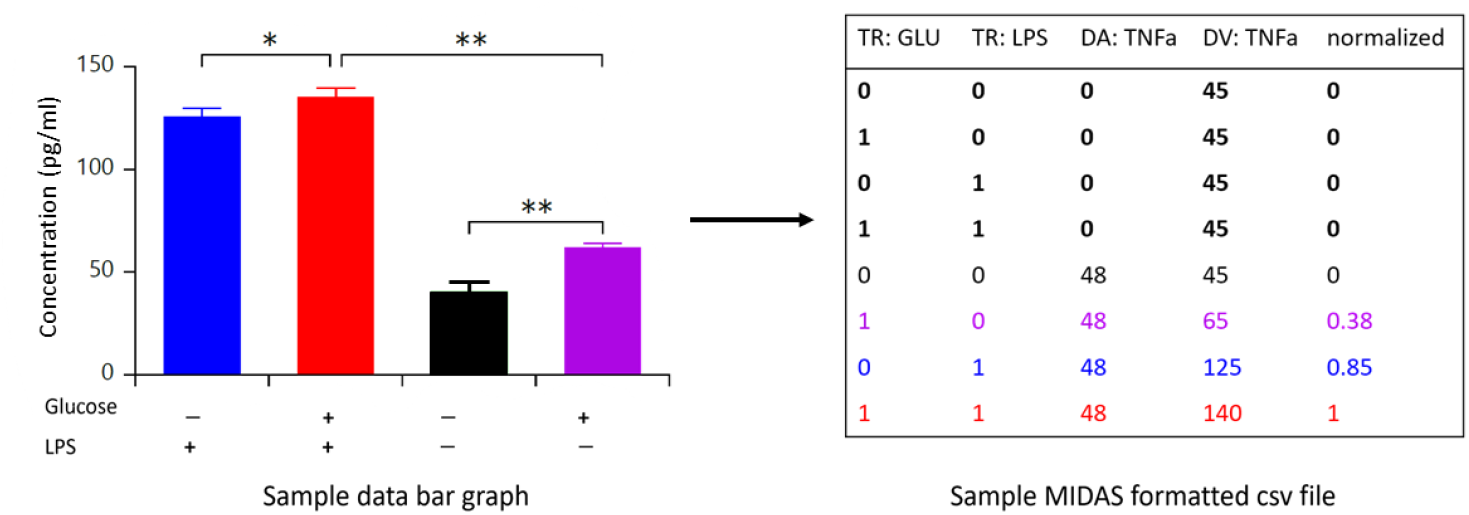
Sample bar graph (left) of protein concentration representing activation from baseline at different glucose (GLU) or lipopolysaccharide (LPS) treatment conditions as found in published studies. The mean values and upper bounds were obtained using WebPlotDigitizer. These values were stored in a MIDAS-formatted comma-separated values (CSV) file, as shown in the table (right). TR:GLU and TR:LPS represent GLU and LPS treatment, respectively, where 0 represents inactive and 1 represents active treatment. DA:TNFa and DV:TNFa represent the time (hour) and concentration (pg/ml) of TNF-α protein, respectively. Normalized values in the table were obtained using the method proposed by Saez-Rodriguez et al. (33). Baseline values (black bar) were used for each treatment condition at the start of the experiment (time 0) in the table (bold text). The values at 48 hours correspond to the respective treatment conditions in the bar graph. For instance, the data obtained from the blue, red, black, and purple bars corresponds to the respective colored rows in the table. TR: treatment. DA: data attribute. DV: data value. TNF: tumor necrosis factor.

Co-culture experimental systems have gained popularity in the field, especially for studying and engineering complex multi-cellular synthetic systems and mimicking tissue structures to some extent (86). We used protein concentrations from individual cell culture studies on either macrophages or endothelial cells for IL-6, TNF-α, VEGF-A, ROS, eNOS, NO, and ROS_ec_ when data from co-culture systems were unavailable (9, 10, 78, 79). We extracted data from figures or tables in these publications (10, 18, 78–81) for calibration and from these publications (9, 29, 81–85) for validation using WebPlotDigitizer, a web-based data extraction software (87). VEGF-A activity under LPS only stimulus was validated using pooled data from (81, 83). VE-cadherin activity under LPS only stimulus was validated using pooled data from (29, 82). We used ImageJ software to quantify Western blot images from published papers when needed. Fig. 3 concisely represents a sample bar graph of protein concentration at different glucose or LPS treatment conditions.

We normalized the curated data for two reasons: (1) to estimate model parameters by fitting model output to data and (2) to pool data from multiple studies. It has been previously shown that it is essential to normalize the data and simulations in the same way to ensure uniformity between model output and data (89). Our model output ranges between 0 and 1. Thus, we normalized the data in this range. Data normalization involves normalizing a data point with respect to the highest value in the series, but this can overemphasize data outliers and underweight minor and recurring differences in the series (33). Instead, we used a multi-step non-linear data normalization technique (33) using the open-source software package CellNOptR (47). To perform data normalization in CellNOptR, the data were first stored as comma-separated values in a MIDAS-formatted file (Fig. 3). Then, the normaliseCNOlist function from the CellNOptR package was used to perform data normalization in R (90). This function performs the normalization using the following steps (illustrated in Fig. S1):

1. The data values are transformed to a fold change for the same experimental treatment condition at time 0.
2. The fold change is transformed using a Hill function similar to *Eq. 2* (91).
3. a penalty for noise in the data (if any) is multiplied by the transformed value to compute the normalized data value.
4. If the Hill-transformed fold changes after noise penalty have negative values, the normalized data are rescaled using the minimum and maximum values in the normalized data list.

We used the no-noise option, which skips step 3. All experimental data values were considered within a large dynamic range [0, ∞), which is the default option for the normaliseCNOlist function. More detailed information on this normalization technique can be found in (33) and CellNOptR reference manual (91).

### Parameter estimation and uncertainty quantification

The LBODEs model was first simulated with default parameters for treatment conditions: *W*_GLU_ = 1 and *W*_LPS_ = 0 for GLU only, *W*_GLU_ = 0 and *W*_LPS_ = 1 for LPS only, and *W*_GLU_ = *W*_LPS_ = 1 for both GLU and LPS. However, the model predictions based on default parameters were only qualitative and not fitted to the data. The model parameters are not available in the literature, and not all parameters have biophysical definitions. Therefore, default parameters were used as a starting point. We performed structural identifiability and observability analyses to minimize uncertainty due to non-identifiable parameters in our model (92, 93). Assessing the model’s structural identifiability and observability helps detect structural issues before model calibration (94, 95). We also performed a global sensitivity analysis using Sobol’s variance-based method (96) to identify the most influential parameters affecting the model response and reduce the parameter space for model calibration. We followed several recommendations from Linden et al. (97) regarding analyses before parameter estimation and best practices for uncertainty quantification. We then conducted parameter estimation on the sensitive parameters. We quantified the uncertainty associated with the estimated parameter values and how this propagates to model prediction uncertainty. Finally, we used local sensitivity analysis on the optimized and validated model as a measure of the effects of local perturbations representing *in silico* knockdowns on model predictions for all species in the PSN. Each of these procedures is detailed in the remainder of this section.

The structural identifiability and observability analyses for the LBODEs model were performed before model calibration using the STRIKE-GOLDD software package in MATLAB (98). The structural identifiability analysis evaluates the possibility of determining the parameter values from output measurements. In contrast, the observability analysis describes the ability to infer the dynamic variables of the model from the model output (94). The structural identifiability is entirely defined by the model equations and input-output mapping and disregards the limitation of data quantity or quality (93–95, 99). The full input-state-parameter observability (FISPO) algorithm in STRIKE-GOLDD was used to perform a joint structural identifiability analysis of model parameters (*W*_*j*_, *n*_*j*_, and 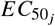 for reactions *j* and τ_*i*_ for species *i*) and observability analysis of all dynamic variables. The observability of the model variables (states, parameters, and inputs) was determined by calculating the rank of a generalized observability-identifiability matrix using Lie derivatives (100). The FISPO analysis yielded a full rank matrix for each computed Lie derivative, which suggests that the LBODEs model is observable and that unknown parameters (*W*_*j*_, *n*_*j*_,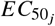, and τ_*i*_) are structurally identifiable when the initial states are known. Further details on using the STRIKE-GOLDD software package and the FISPO algorithm are available elsewhere (94, 98, 100).

We used Sobol’s variance-based method to perform a global sensitivity analysis of the model parameters (96). The UQLab toolbox (101) was used to perform the global sensitivity analysis (102) in MATLAB. We assessed the effect of reaction weight (*W*_*j*_), Hill coefficient (*n*_*j*_), half effect 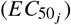,and time constant (τ_*i*_) on the model response. First, we sampled the parameter from ranges specified in a previous study on uncertainty quantification of an LBODEs model also developed using Netflux (76). For this analysis, *W*_*j*_, *n*_*j*_, 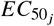,and τ_*i*_ were constrained in the ranges of [0.8, 1], [1.4, 4], [0.4, 0.6], and [0.5, 10], respectively. A Monte Carlo estimator in the UQLab sensitivity analysis toolbox was used to sample from the parameters in respective ranges. The sensitivity indices were computed using Sobol pseudo-random sampling (101, 102). The sample size for Monte Carlo estimation was set to 1000. The sample size was chosen based on the optimum computation time for a global sensitivity analysis on 144 parameters. The Sobol sensitivity analysis computes sensitivity indices for scalar output quantities (97). We chose the predicted responses of 8 of the 10 output nodes in Fig. 1b (IL-6, TNF-α, IL-1β, VEGF-A, ROS, eNOS, NO, and ROS_ec_) at 48 hours as our scalar quantities of interest. We computed and plotted the first-order (Supplemental Fig. S2) and total Sobol sensitivity indices (Supplemental Fig. S3) for the quantities of interest organized by each type of parameter. We selected the most sensitive parameters as those that had a total Sobol sensitivity index above a given threshold or cut-off value. Identifying a single cut-off absolute value for all the parameters (*W*_*j*_, *n*_*j*_, 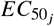,and τ_*i*_) was challenging. Therefore, we assumed a different threshold value for each parameter set as in Linden et al. (97). The total order Sobol sensitivity index within 10% of the maximum sensitivity index was fixed as the threshold for each parameter set. A total of 82 parameters (from the 204 total for the LBODEs model: the three reaction parameters *W*_*j*_, *n*_*j*_, and 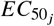 for the 38 downstream reactions in Table 2 and the one species parameter τ_*i*_ for each of the three treatment conditions for the 30 species in Table 3) were identified as sensitive and were marked for parameter estimation. The non-sensitive parameters were set to their default values. More information on using the UQLab toolbox for global sensitivity analysis can be found in the user manual (102).

We estimated the sensitive parameters by calibrating the LBODEs model to the normalized curated datasets for each treatment condition. The sensitive parameters were optimized using a multi-start non-linear least squares parameter optimization method to avoid converging to a local minimum (103). We sampled 100 sets of parameter start values using the Latin hypercube sampling technique (104). Each parameter was sampled uniformly across the following ranges: [0.9, 1], [1.4, 4], [0.001, 0.84], and [0.01, 10] for *W*_*j*_, *n*_*j*_, 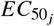,and τ_*i*_, respectively. The Latin hypercube sampling technique combined these parameter samples to cover the parameter space efficiently. We used the Latin hypercube sampling routine in MATLAB as implemented in functions provided online (105) using published methods (104, 106). Each of the sampled sets of parameter start values was used as initial guesses in the fmincon nonlinear programming solver in MATLAB, which returns an optimized parameter set.

`fmincon` minimized the objective function, which was the sum of squared error (SSE) between model predictions and published data (Fig. 3). The SSE was computed between observed responses from data at available time points and model-predicted responses for the following output nodes in Fig. 1b: IL-6, TNF-α, IL-1β, VEGF-A_mRNA_, ROS, eNOS, NO, and ROS_ec_. First, we performed a multi-start optimization of sensitive *W*_*j*_, *n*_*j*_, 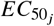,and τ_*i*_ parameters for the GLU only treatment condition, where *W*_GLU_ = 1 and *W*_LPS_ = 0. For the GLU only condition, initial values 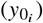 of VEGF-A, NO, and eNOS were set to 1 to depict relative downregulation. It is relevant to note that the parameter optimization was carried out sequentially in the order of occurrence of species nodes from top to bottom in the network to avoid bias from multiple data points. For the GLU only condition, model parameters governing the expression of ROS, ROS_ec_, NO, eNOS, and VEGF-A were fitted against data before fitting the parameters for IL-6, TNF-α, and IL-1β response. The optimized reaction parameters (*W*_*j*_, *n*_*j*_, and 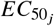) were fixed across different treatment conditions (GLU only, LPS only, both GLU and LPS). The sensitive τ_*i*_ parameters were re-calibrated for the LPS only and both GLU and LPS treatment conditions. The re-calibration of τ_*i*_ values was necessary for different treatment conditions as the strength of the input signal affects rapid or delayed species response. The multi-start parameter estimation procedure generated 100 sets of fitted parameters for each treatment condition and an optimal (best) parameter set that gave the smallest SSE between model predictions and data. From the 100 sets of fitted parameters for each treatment condition, a set was labeled as part of the acceptable subset of parameters if the SSE for the set was within 20% of the smallest SSE. The means of the parameters within the acceptable subset were defined as the nominal parameter set for each condition. These nominal parameters are reported in Tables 2 and 3. Estimated values of sensitive parameters are displayed with three significant figures to distinguish them from the non-sensitive parameters at their default values.

We quantified the uncertainty associated with model predictions using Monte Carlo ensemble simulation, also known as sampling-based uncertainty propagation (97, 107). This approach repeatedly simulates a model using samples from the posterior distributions of the parameters as inputs to obtain a mapping to results (107), producing a posterior distribution of model predictions for each treatment case. To obtain a measure of the uncertainty of the parameters, we used the function randsample(population, numSamples) in MATLAB to return numSamples values sampled uniformly at random from the values in the vector population. With numSamples at 5000, we called randsample independently for each sensitive parameter. For each parameter, we specified population as the values of that parameter in the acceptable parameters subset for each treatment condition that resulted from the multi-start parameter optimization approach described in the previous paragraph. The resulting 5000 values for each parameter were considered as the posterior distribution for the parameter. The posterior distributions for the parameters are shown along with the nominal parameter values (means of acceptable parameters subset) in Supplemental Fig. S4. We stored the values returned from randsample for all the parameters by their sample number. For a given sample number, each parameter was a sample from its posterior distribution, independent of the other parameter values. Together, these parameter values at the same sample number comprised a set of parameters that were run through the LBODEs model to obtain model predictions. The Monte Carlo ensemble simulation involved repeatedly running the LBODEs model for each of the 5000 combinations of parameter sets organized by sample number. The resulting posterior distributions of model predictions were used to calculate the 95% equal-tail credible interval at each time point for each response. The 95% equal-tail credible interval is bound by a lower limit representing the 2.5^th^ percentile of the posterior distribution and upper limit representing the 97.5^th^ percentile of the posterior distribution (108). In other words, a 95% equal-tail credible interval suggests that there is a 95% probability that a true estimate is within this interval, and all values inside the interval are more likely to represent the parameter than the values outside the interval (108, 109). The use of 95% equal-tail credible intervals provides better interpretability of these estimates. Showing model results with credible intervals from posterior distributions of model predictions has been widely adopted in systems biology (97, 99, 109, 110). The statistical analysis of the posterior distribution of parameters and model predictions through credible intervals and visual checks allowed us to understand the extent of uncertainty in the estimated parameters (Supplemental Figs. S4–S6).

The robustness of reaction rules and species in the model to perturbations for plausible *in silico* knockdowns were analyzed using a local sensitivity analysis with finite difference approximation:

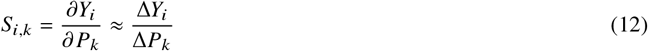

where *S*_*i*,*k*_ is a sensitivity coefficient for a given output species *i* and parameter *k, Y*_*i*_ (*P*) is the output for species *i* at the nominal parameter values *P*, Δ*Y*_*i*_ = *Y*_*i*_ (*P* +Δ*P*_*k*_) − *Y*_*i*_ (*P*) is the change in output species *i* calculated at the perturbed parameter value, and Δ*P*_*k*_ is a small perturbation in parameter *k*.

## RESULTS

### Model calibration

The simulation results are shown in Figs. 4–6 for the LBODEs model calibrated to the normalized biochemical data for three different treatment conditions, (1) active GLU stimulus, (2) active LPS stimulus, and (3) active GLU and LPS stimuli, respectively. Based on the published co-culture experimental studies, the model was simulated for short-term exposure (48 hours). For each treatment condition, most fitted simulations for each output response captured the observed time-dependent data very well. The posterior distributions and means of acceptable reaction parameters (*W*_*j*_, *n*_*j*_, 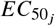) are plotted at 12 hours for respective reaction index *j* and rule as reported in Table 2 in Supplemental Fig. S4. Supplemental Fig. S5 shows the posterior distributions and means of acceptable time constant (τ_*i*_) parameter values for different treatment conditions at 12 hours. Further, the distributions of prediction posteriors for IL-6, TNF-α, IL-1β, VEGF-A_mRNA_, ROS, eNOS, NO, ROS_ec_, VEGF-A, and Gap Width at 12 hours for each condition are reported in Supplemental Fig. S6. The distribution of predicted responses captures the prediction uncertainty at 12 hours for the respective output species. We observed a low prediction uncertainty in most output responses, as demonstrated by narrow 95% credible intervals. Also, most observed data were within the 95% credible intervals of the model predictions.

**Figure 4:**
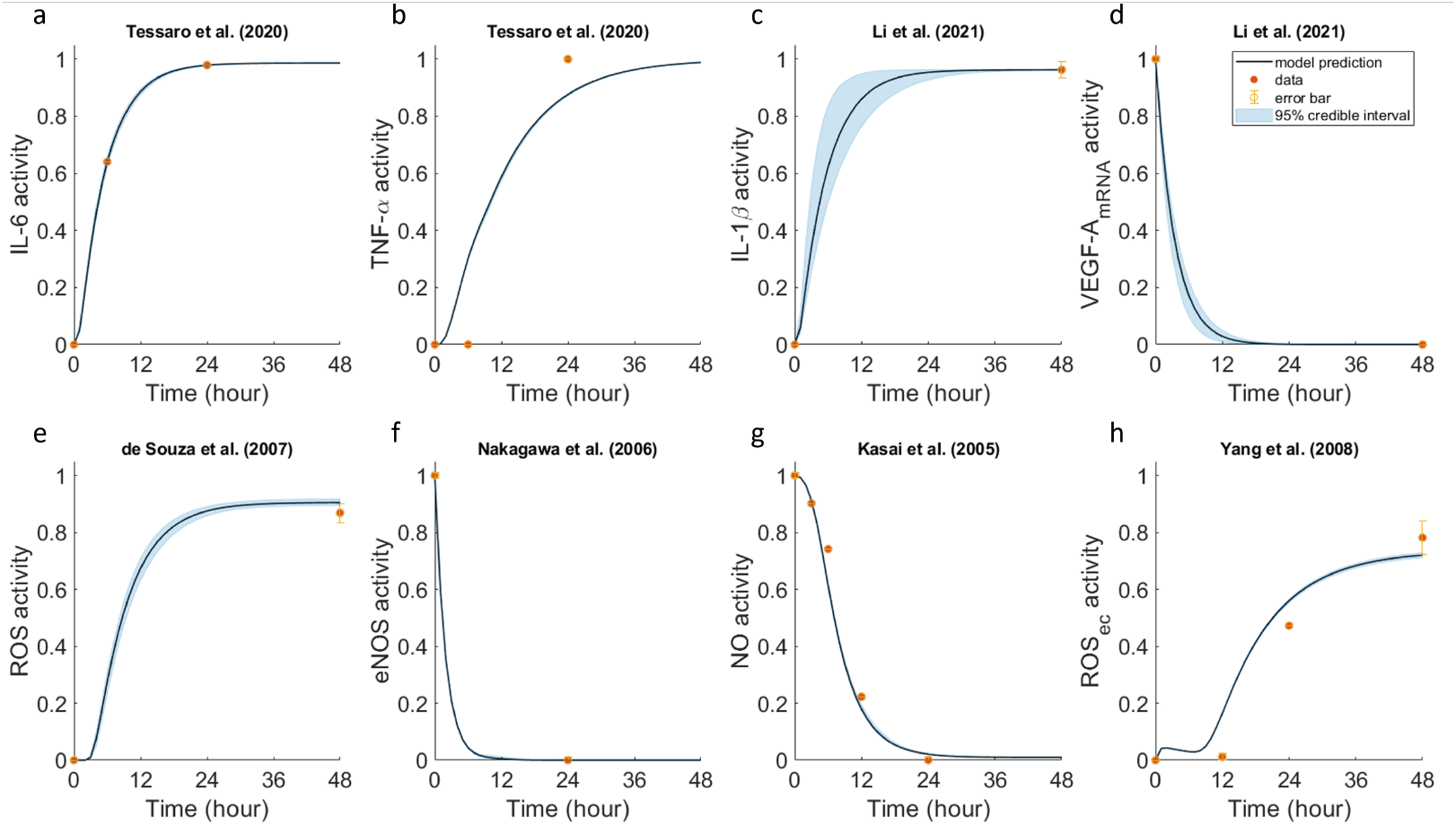
PSN responses to glucose (GLU) for 48 hours (*W*_GLU_ = 1 and *W*_LPS_ = 0) for calibration to biochemical data available from cell culture experiments in Tessaro et al. (10), Li et al. (18), de Souza et al. (78), Nakagawa et al. (79), Kasai et al. (80), and Yang et al. (81). Dynamic LBODEs model predicted responses are compared to normalized protein levels of (a) IL-6, (b) TNF-α, (c) IL-1β, (d) VEGF-A_mRNA_ levels, (e) ROS fluorescence, (f) eNOS, (g) NO, and (h) ROS_ec_. The model predictions at nominal parameter values are shown as black solid curves, data are shown as orange points, and the 95% credible intervals for the calibration are shown as blue-shaded regions. The subfigure titles provide the sources for the data in the corresponding plots. The subscript “ec” denotes intracellular expression in glomerular endothelial cells. Abbreviations are defined in Table 1.

Under fixed GLU treatment, the upregulated pro-inflammatory cytokines (IL-6, TNF-α, IL-1β) and ROS activity reached a steady state around 24 hours. As the responses showed the widest credible intervals around 12 hours (Fig. 4c–e), the posterior distributions of the prediction responses for the 10 output nodes in Fig. 1b are shown at 12 hours (Supplemental Fig. S6). The fitted IL-1β response had relatively high prediction uncertainty, as observed from the wide 95% credible interval (Fig. 4c). More experimental evidence at different time points could reduce this prediction uncertainty. Glucose-mediated downregulation of VEGF-A_mRNA_ (Fig. 4d) caused the decrease in eNOS and NO activity (Fig. 4f–g). Although the PSN model considers a simplified VEGF-dependent eNOS/NO pathway, it efficiently captured the glucose-mediated NO downregulation. Under high glucose conditions, we identified that the degree of influence of Ca on NO, i.e., the reaction weight for reaction rule Ca ⇒ NO (*j* = 40), also contributed to 100% downregulation of NO activity.

Calibrating time constant parameters for each treatment condition separately (Table 3) allowed us to capture the earlier activation of IL-6, TNF-α, and IL-1β cytokines for LPS treatment (Fig. 5a–c). LPS-mediated output responses achieved 50% of maximum activation as early as 6 hours when compared to GLU only treatment, which demonstrates the difference between the strength of glucose (Fig. 4) and LPS (Fig. 5) stimuli. Under LPS treatment, a relatively higher ROS_*ec*_ activity was observed (Fig. 5h compared to Fig. 4h) and mainly mediated by NO uncoupling from its pathway. The reaction weight of eNOS ⇒ ROS_*ec*_ (*j* = 34) was relatively higher than for NADPH_*ec*_ ⇒ ROS_*ec*_ (*j* = 25) (Table 2). LPS-mediated VEGF-A_mRNA_ upregulation (Fig. 5d) initiated the increase in eNOS and NO activity (Fig. 5f–g). ROS expression was upregulated due to the PI3K-dependent pathway in macrophages (Fig. 5e) and the eNOS-dependent pathway in GECs (Fig. 5h). Also, note that the fitted IL-1β response had smaller prediction uncertainty (narrow 95% credible interval) compared to the GLU only case due to an additional data point for the LPS only case (Fig. 5c compared to Fig. 4c).

**Figure 5:**
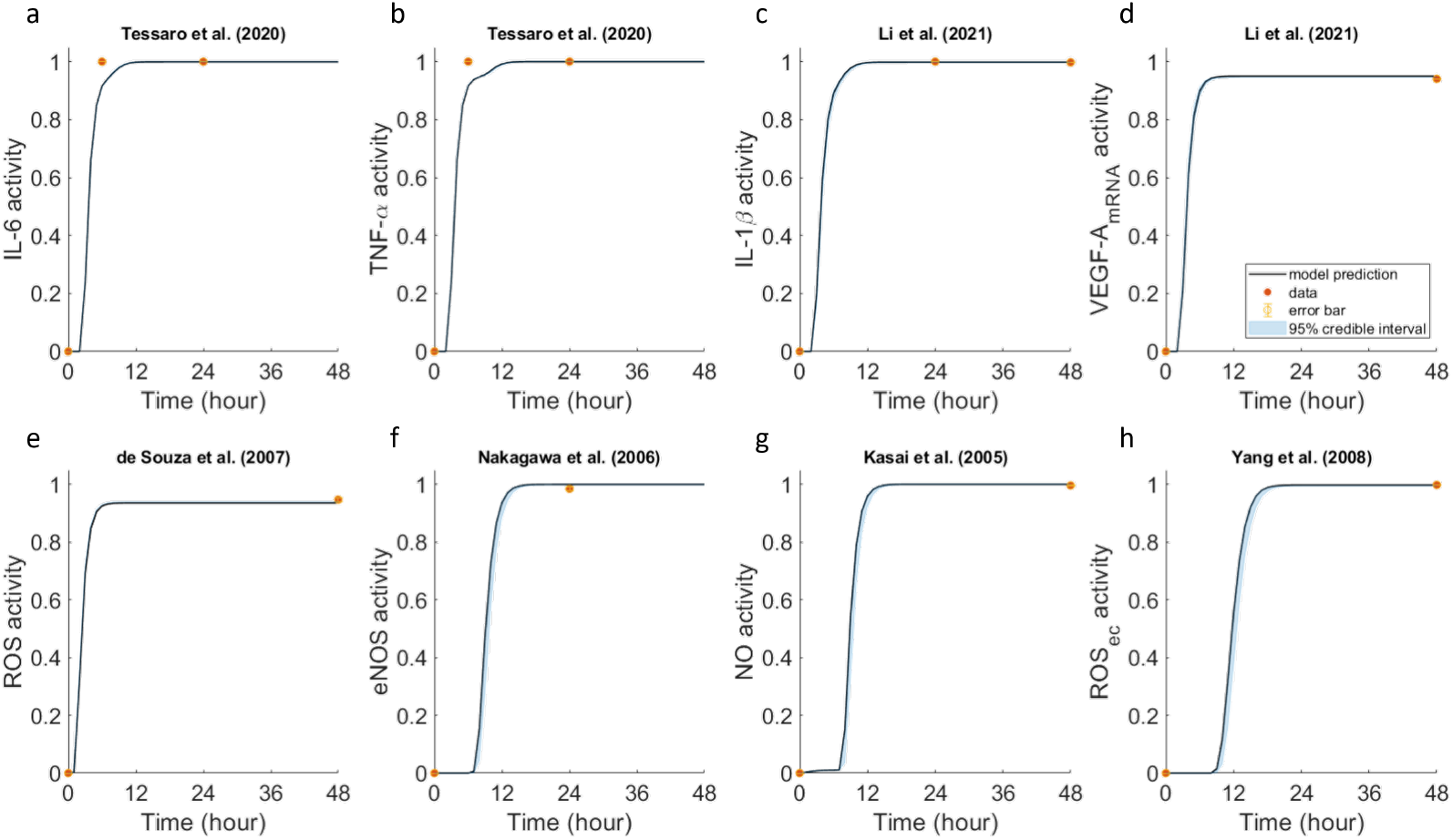
PSN responses to lipopolysaccharide (LPS) for 48 hours (*W*_GLU_ = 0 and *W*_LPS_ = 1) for calibration to biochemical data available from cell culture experiments in Tessaro et al. (10), Li et al. (18), de Souza et al. (78), Nakagawa et al. (79), Kasai et al. (80), and Yang et al. (81). Dynamic LBODEs model predicted responses are compared to normalized protein levels of (a) IL-6, (b) TNF-α, (c) IL-1β, (d) VEGF-A_mRNA_ levels, (e) ROS fluorescence, (f) eNOS, (g) NO, and (h) ROS_ec_. The model predictions at nominal parameter values are shown as black solid curves, data are shown as orange points, and the 95% credible intervals for the calibration are shown as blue-shaded regions. The subfigure titles provide the sources for the data in the corresponding plots. The subscript “ec” denotes intracellular expression in glomerular endothelial cells. Abbreviations are defined in Table 1.

When both GLU and LPS stimuli were active, LPS stimulus dominated the expression of growth factors, cytokines, and ROS, as observed from their early activation, except for IL-1β (Fig. 6c), NO (Fig. 6g), and ROS_*ec*_ (Fig. 6h). The fitted VEGF-A_mRNA_ activity (Fig. 6d) increased after an initial delay compared to the LPS only case, possibly due to the combined effects countering that of GLU only (decrease from baseline). NO upregulation (Fig. 6g) was also delayed due to the combined negative effect of GLU and the positive effect of LPS on NO. The fitted IL-1β and NO response had the highest uncertainty in estimates as observed from the wide 95% credible intervals (Fig. 6c,g), which could be resolved with more comprehensive experimental data. The broad posterior distribution of uncertainty in the time constant value for NO (Supplemental Fig. S5) is the source for prediction uncertainty or wide credible intervals. The posterior distributions of predictions for IL-1β and NO with both GLU and LPS showed a wide range of predicted values at 12 hours, unlike other species (Supplemental Fig. S6).

**Figure 6:**
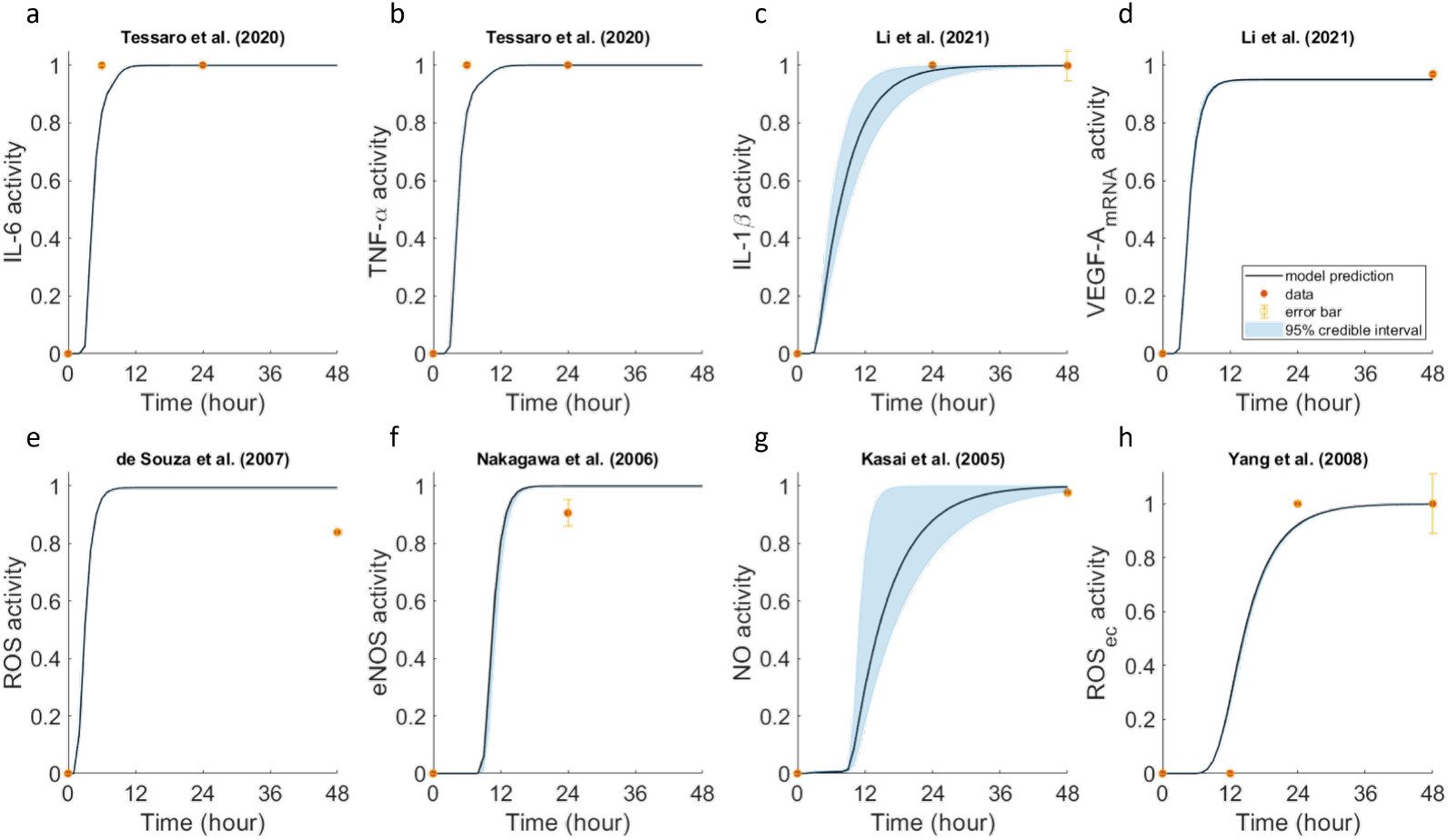
PSN responses to both glucose (GLU) and lipopolysaccharide (LPS) for 48 hours (*W*_GLU_ = 1 and *W*_LPS_ = 1) for calibration to biochemical data available from cell culture experiments in Tessaro et al. (10), Li et al. (18), de Souza et al. (78), Nakagawa et al. (79), Kasai et al. (80), and Yang et al. (81). Dynamic LBODEs model predicted responses are compared to normalized protein levels of (a) IL-6, (b) TNF-α, (c) IL-1β, (d) VEGF-A_mRNA_ levels, (e) ROS fluorescence, (f) eNOS, (g) NO, and (h) ROS_ec_. The model predictions at nominal parameter values are shown as black solid curves, data are shown as orange points, and the 95% credible intervals for the calibration are shown as blue-shaded regions. The subfigure titles provide the sources for the data in the corresponding plots. The subscript “ec” denotes intracellular expression in glomerular endothelial cells. Abbreviations are defined in Table 1.

### Model validation

The predicted activity for output responses compared well against most of the validation data not used in the calibration (Figs. 7–9). For GLU only, the predicted ROS_*ec*_ activity did not capture the validation dataset well. Although VEGF-A activity was not fitted to experimental data (instead, VEGF-A (mRA) activity was fitted), the predicted VEGF-A activity agrees with the validation dataset for GLU treatment (Fig. 7d). The predicted VEGF-A activity for the other treatment conditions did not match as well with observed data (Fig. 8d,Fig. 9d) with a consistent offset. The estimated *W*_*j*_ for VEGF-A_mRNA_ ⇒ VEGF-A protein conversion (*j* = 17) was less than 1 (Table 2), which explains why the VEGF-A activity reaches a steady state below 1. Due to the lack of quantitative data for intercellular gap width, we validated the pJunction node against data for the expression of phosphorylated VE-cadherin, a type of junction protein, instead (Fig. 8e). The change in intercellular Gap Width in the PSN depends on the phosphorylation of cell-cell junction proteins such as VE-cadherin, which results in the weakening of endothelial junction integrity (24, 111). The predicted phosphorylated VE-cadherin activity (pJunction node) agreed quite well with the validation data in the LPS only case (Fig. 8e).

**Figure 7:**
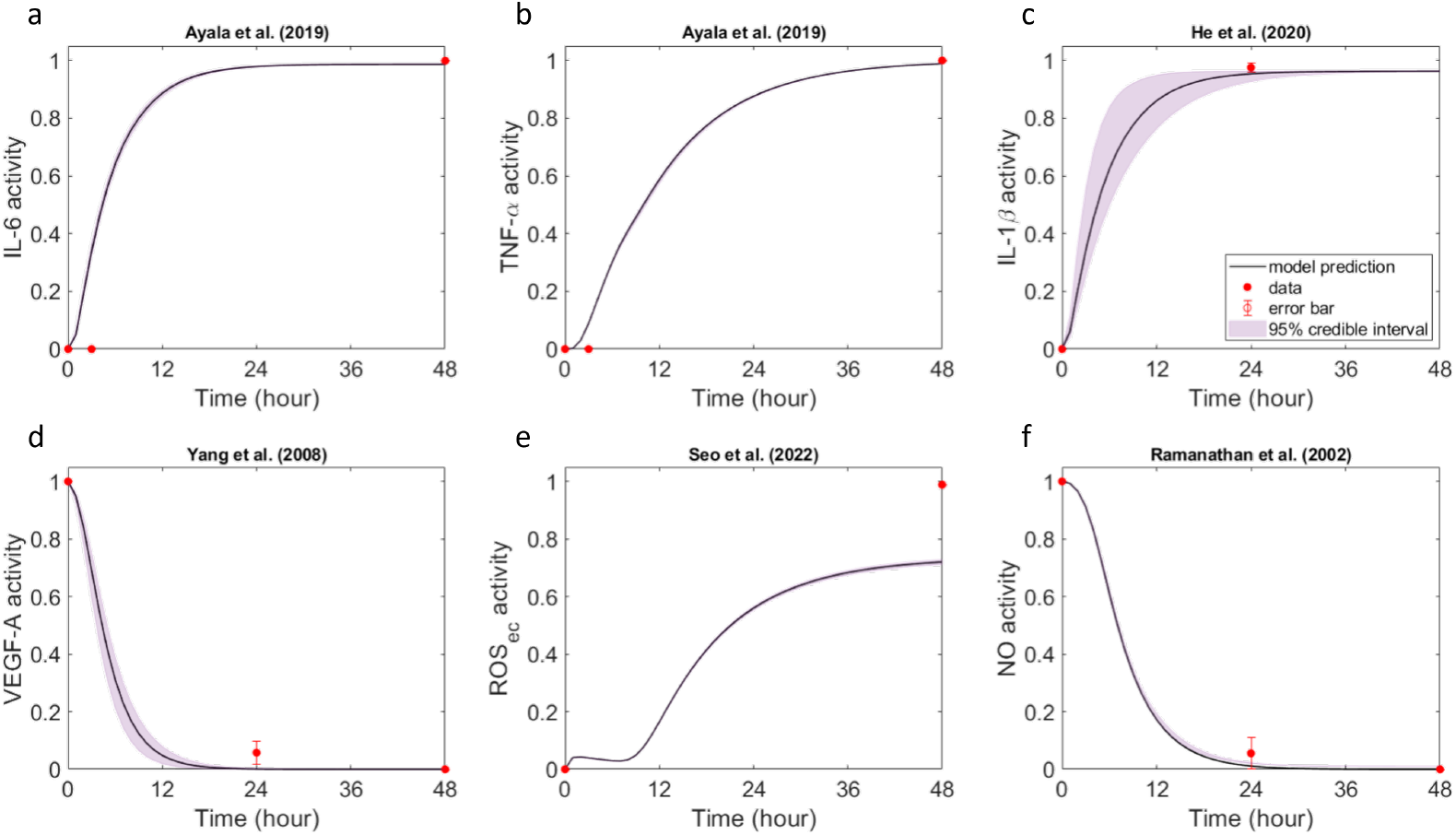
PSN responses to glucose (GLU) for 48 hours (*W*_GLU_ = 1 and *W*_LPS_ = 0) for validation against biochemical data available from cell culture experiments in Ayala et al. (9), He et al. (84), Yang et al. (81), Seo et al. (29), and Ramanathan et al. (85). Dynamic LBODEs model predicted responses are compared for selected species against available normalized protein levels of (a) IL-6, (b) TNF-α, (c) IL-1β protein levels, (d) VEGF-A levels, (e) ROS_ec_, and (f) NO. The model predictions at nominal parameter values are shown as black solid curves, data are shown as orange points, and the 95% credible intervals for the validation are shown as purple-shaded regions. The subfigure titles provide the sources for the data in the corresponding plots. The subscript “ec” denotes intracellular expression in glomerular endothelial cells. Abbreviations are defined in Table 1.

**Figure 8:**
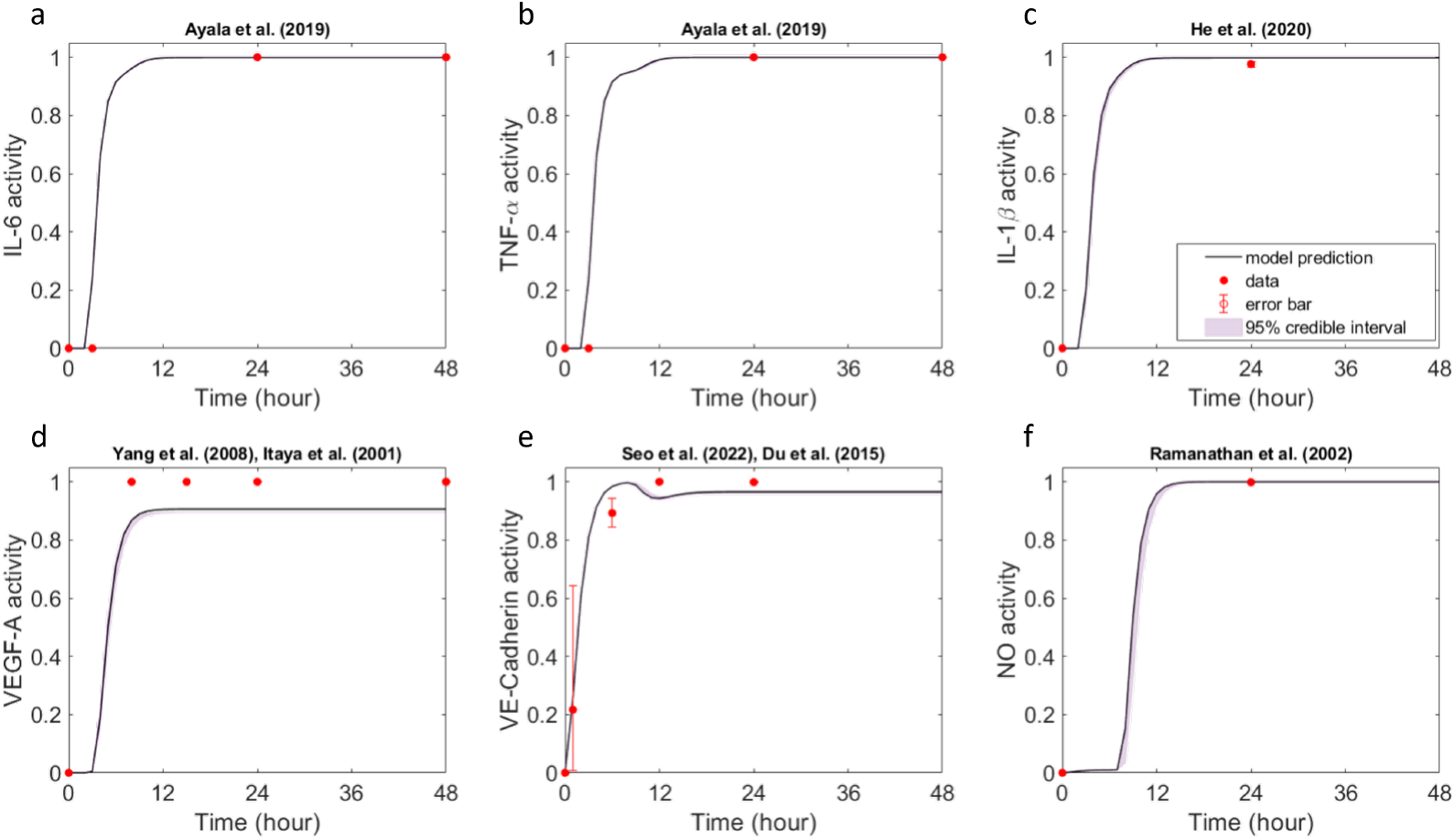
PSN responses to lipopolysaccharide (LPS) for 48 hours (*W*_GLU_ = 0 and *W*_LPS_ = 1) for validation against biochemical data available from cell culture experiments in Ayala et al. (9), He et al. (84), Yang et al. (81), Itaya et al. (83), Seo et al. (29), Du et al. (82), and Ramanathan et al. (85) upon stimulation with lipopolysaccharide (LPS) for 48 hours (*W*_GLU_ = 0 and *W*_LPS_ = 1). Dynamic LBODEs model predicted responses are compared for selected species against available normalized protein levels of (a) IL-6, (b) TNF-α, (c) IL-1β protein levels, (d) VEGF-A levels, (e) phosphorylated VE-cadherin (representing the pJunction node in the network), and (f) NO. The model predictions at nominal parameter values are shown as black solid curves, data are shown as orange points, and the 95% credible intervals for the validation are shown as purple-shaded regions. The subfigure titles provide the sources for the data in the corresponding plots. Abbreviations are defined in Table 1.

**Figure 9:**
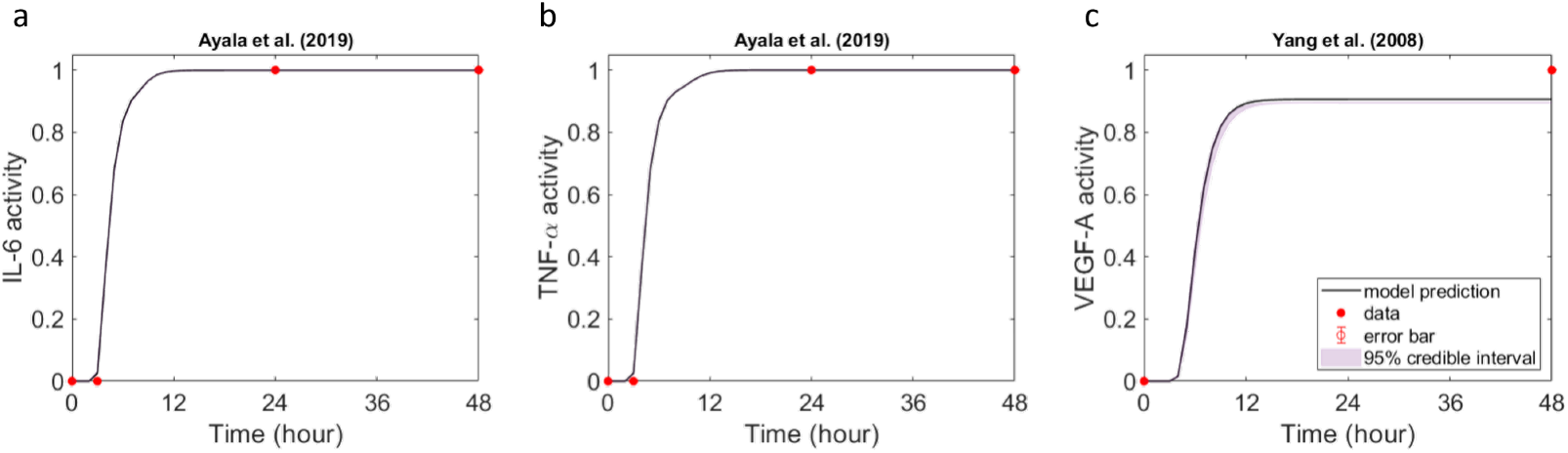
PSN responses to both glucose (GLU) and lipopolysaccharide (LPS) for 48 hours (*W*_GLU_ = 1 and *W*_LPS_ = 1) for validation against biochemical data available from cell culture experiments in Ayala et al. (9) and Yang et al. (81) upon stimulation with both glucose (GLU) and lipopolysaccharide (LPS) for 48 hours (*W*_GLU_ = 1 and *W*_LPS_ = 1). Dynamic LBODEs model predicted responses are compared for selected species against available normalized protein levels of (a) IL-6, (b) TNF-α, and (c) VEGF-A. The model predictions at nominal parameter values are shown as black solid curves, data are shown as orange points, and the 95% credible intervals for the validation are shown as purple-shaded regions. The subfigure titles provide the sources for the data in the corresponding plots. Abbreviations are defined in Table 1.

### PSN regulatory species dynamics

We plotted the dynamics of the regulatory species in the PSN that were not used in model fitting (white regulatory nodes in Fig. 1b) for the three different treatment conditions (Figs. 10–12). The species dynamics show the behaviors of signaling molecules, transcription factors, and receptors for which dynamic experimental data that met our inclusion criteria were unavailable. The model predicted a short-term upregulation of VEGFR1, VEGFR2, PLC-γ, PI3K_*ec*_, and AKT_*ec*_ that returned to baseline (Fig. 10d) following VEGF-A downregulation under GLU treatment. The model predicted that an increase in Gap Width was directly related to Ca and pJunction activity. In the absence of GLU treatment, the predicted activity of AGE, RAGE, RAGE_*ec*_, NADPH, and NADPH_*ec*_ remained at baseline (Fig. 11a,e).

**Figure 10:**
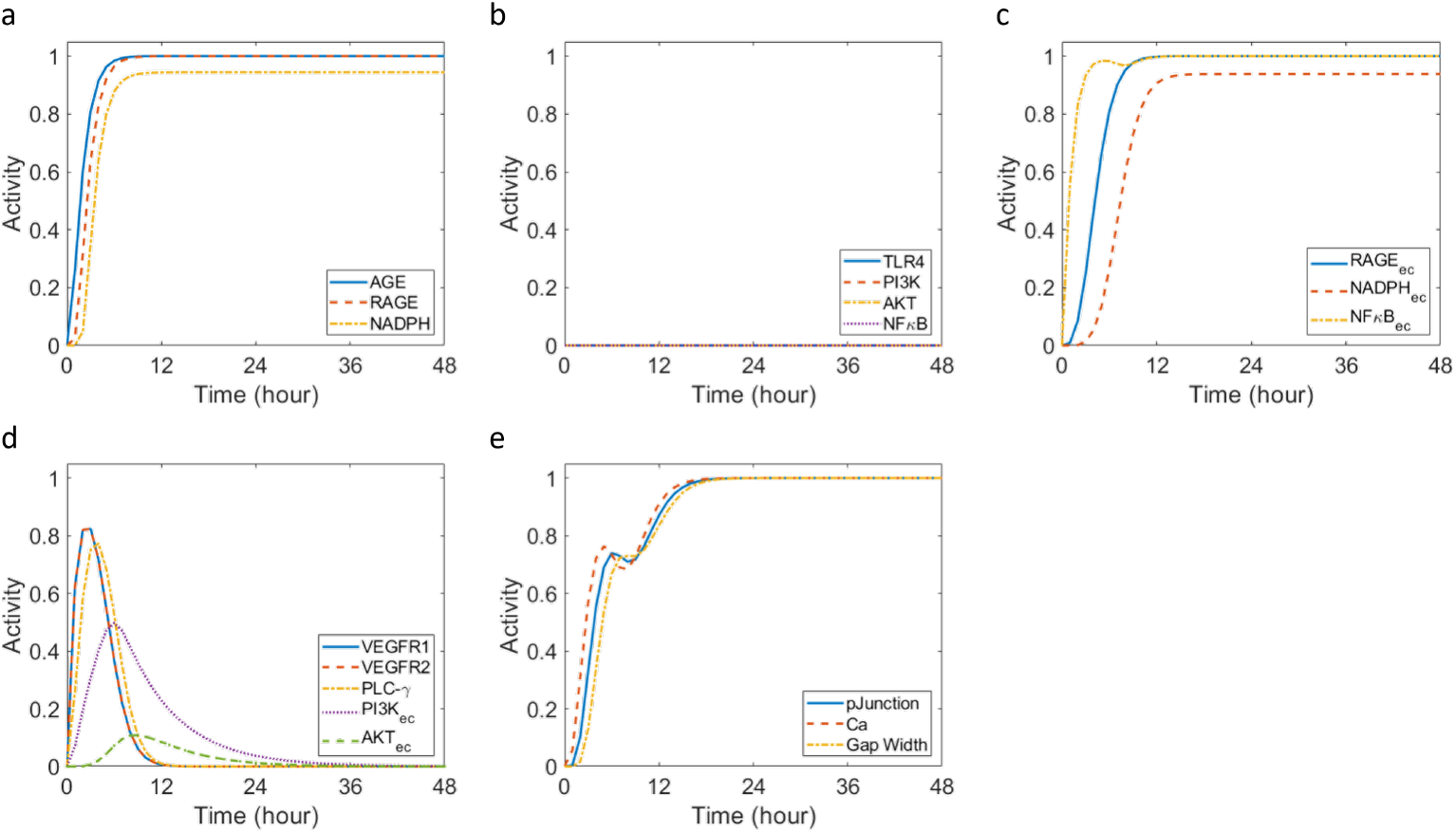
PSN responses to glucose (GLU) for 48 hours (*W*_GLU_ = 1 and *W*_LPS_ = 0) showing dynamic LBODEs model predictions for all regulatory species at nominal parameter values. (a) Regulatory species activity associated with GLU only stimulation. (b) Regulatory species activity associated with LPS only stimulation. (c) Regulatory species activity observed in glomerular endothelial cells. (d) Regulatory species activity associated with VEGF-A activation. (e) Regulatory species activity associated with Gap Width change. The subscript “ec” denotes intracellular expression in the glomerular endothelial cells. Abbreviations are defined in Table 1.

**Figure 11:**
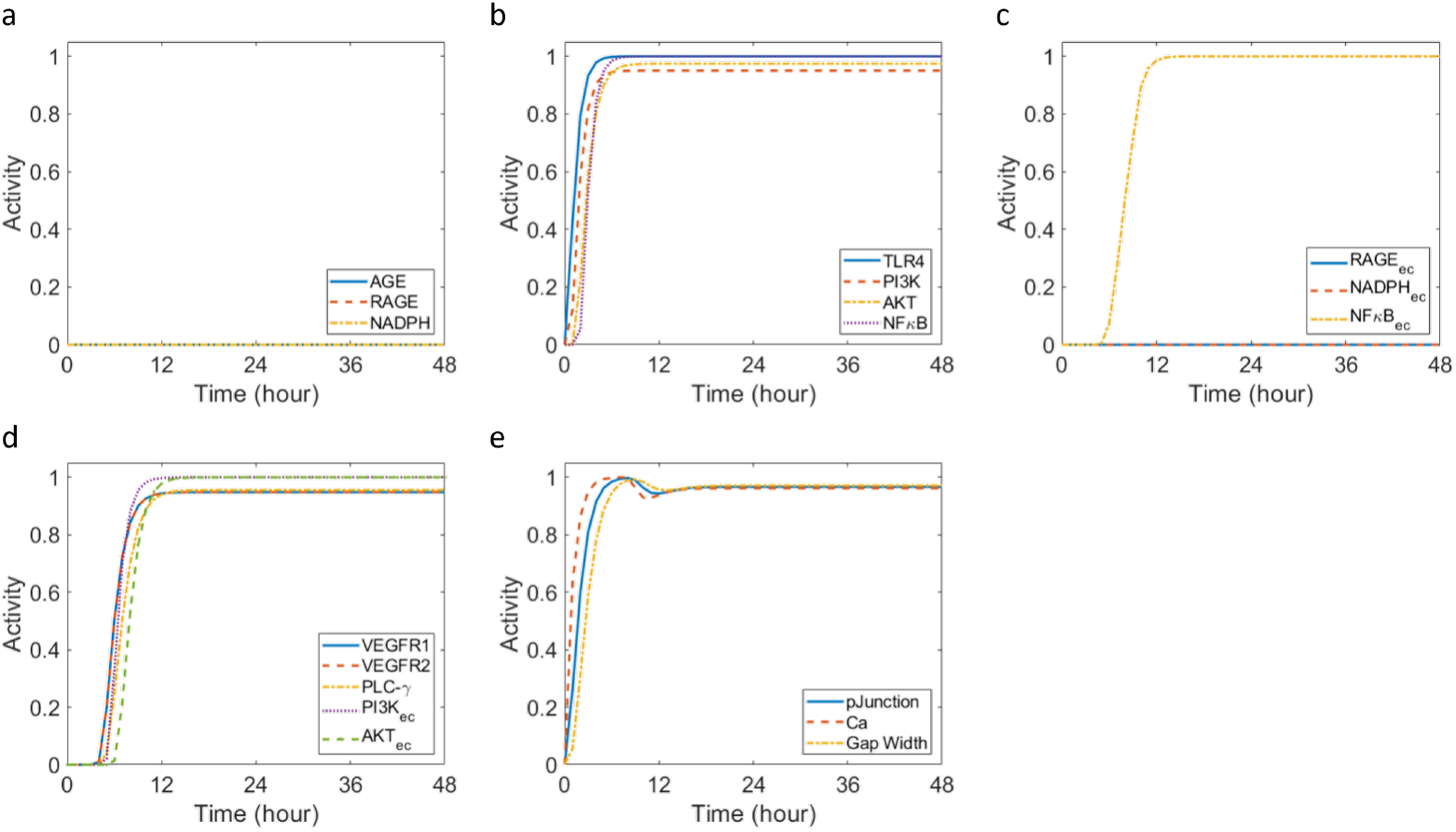
PSN responses to lipopolysaccharide (LPS) for 48 hours (*W*_GLU_ = 0 and *W*_LPS_ = 1) showing dynamic LBODEs model predictions for all regulatory species at nominal parameter values. (a) Regulatory species activity associated with GLU only stimulation. (b) Regulatory species activity associated with LPS only stimulation. (c) Regulatory species activity observed in glomerular endothelial cells. (d) Regulatory species activity associated with VEGF-A activation. (e) Regulatory species activity associated with Gap Width change. The subscript “ec” denotes intracellular expression within glomerular endothelial cells in the network. Abbreviations are defined in Table 1.

The model does not consider the competitive binding of VEGF-A to VEGFR1- and VEGFR2 receptors (Fig. 12d). This was based on the calculated total order sensitivities (Supplemental Fig. S3) for the following reaction rules that were relatively insensitive: VEGF-A VEGFR1 (*j* = 19), VEGF-A ⇒ VEGFR2 (*j* = 20), VEGFR1 ⇒ PI3K_*ec*_ (*j* = 24), and VEGFR2 ⇒ PI3K_*ec*_ (*j* = 23). The strength of these interactions (*W*_*j*_) was equal to 1 (Table 2). However, we consider different VEGFR1- and VEGFR2-mediated downstream signaling in the PSN. VEGFR1 promotes downstream pro-inflammatory signaling via the PLC-γ pathway, whereas VEGFR2 promotes higher kinase activity via PI3K/AKT activation. We also observed that glucose promoted PI3K_*ec*_ activity for longer (24 hours) when compared to PLC-γ activity (12 hours) (Fig. 10d). Under combined treatment, PI3K_*ec*_ kinase activity also preceded PLC-γ activity (Fig. 12d).

**Figure 12:**
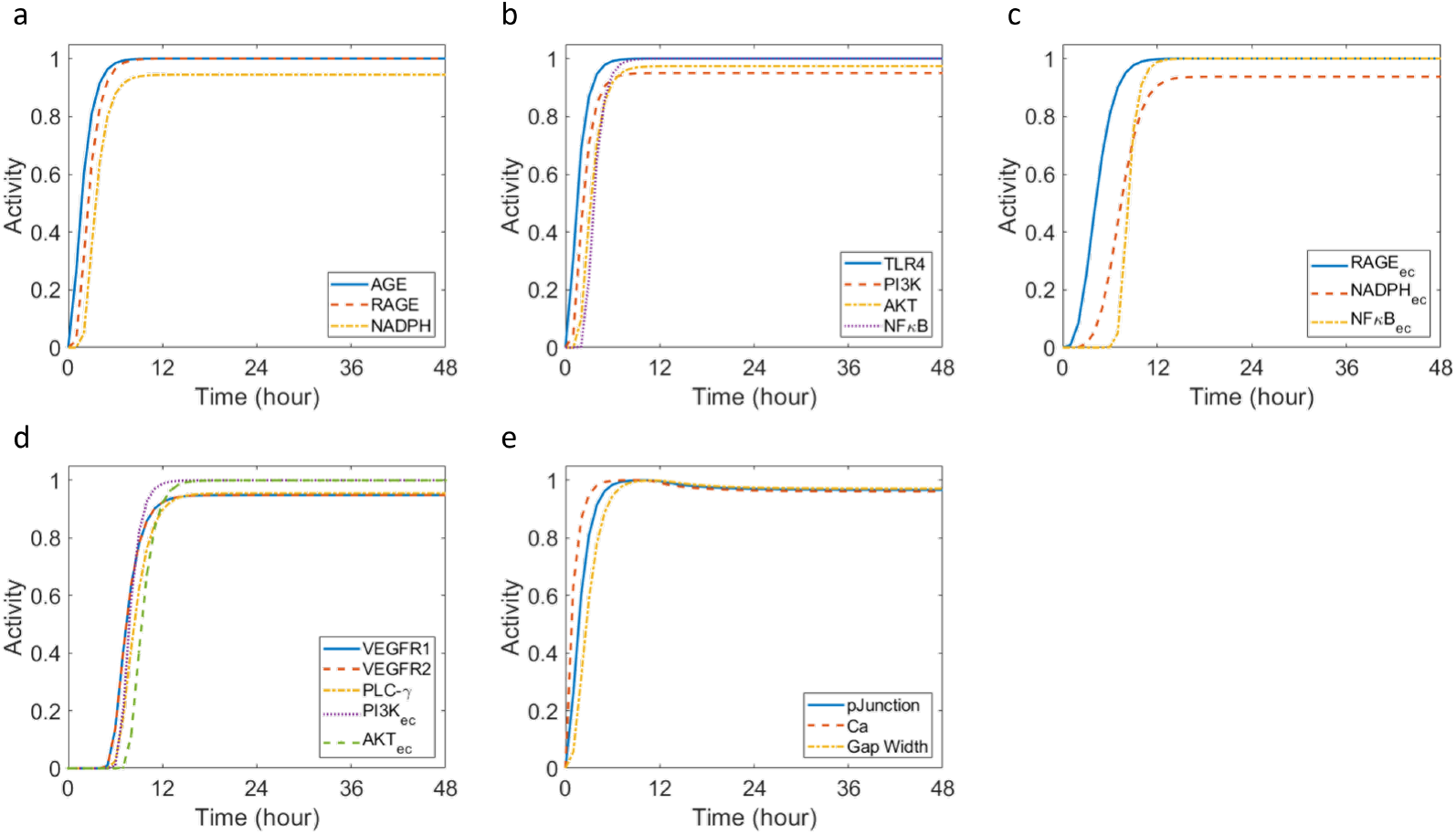
PSN responses to both glucose (GLU) and lipopolysaccharide (LPS) for 48 hours (*W*_GLU_ = 1 and *W*_LPS_ = 1) showing dynamic LBODEs model predictions for all regulatory species at nominal parameter values. (a) Regulatory species activity associated with GLU only stimulation. (b) Regulatory species activity associated with LPS only stimulation. (c) Regulatory species activity observed in glomerular endothelial cells. (d) Regulatory species activity associated with VEGF-A activation. (e) Regulatory species activity associated with Gap Width change. The subscript “ec” denotes intracellular expression within glomerular endothelial cells in the network. Abbreviations are defined in Table 1.

### Model predictions under partial activation or knockdown

The calibrated and validated LBODEs model was used to make predictions under other combinations of stimuli beyond those tested experimentally in the literature. The model simulations shown previously were performed with combinations of fully active stimuli. Here, we predicted output responses under partially active stimuli. To modulate the strength of GLU and LPS stimuli, we varied the reaction weights (*W*_*j*_) for GLU and LPS. The other reaction parameters are set at nominal values unless otherwise indicated (Table 2). We used *Eq. 12* to analyze the robustness of reaction rules and species in the model to perturbations for plausible *in silico* knockdowns, specifically in response to one-at-a-time changes in *W*_*j*_ for all reactions and 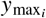 for all species.

First, we simulated the effects of a partially active GLU stimulus (0≤ *W*_GLU_≤ 1) to explore ranges of hyperglycemia in the early stage of DKD while LPS was not active (*W*_LPS_ = 0) mimicking a non-inflammatory state (Fig. 13). We considered the nominal species parameters for this analysis to be those for GLU only treatment (Table 3). We observed that 50% of the maximum glucose activity (*W*_GLU_ ≥0.25) was sufficient to activate and express IL-6 cytokine (Fig. 13a). Low glucose activity was insufficient for maintaining upregulated IL-6 levels, and IL-6 activity returned to the baseline unless GLU was fully activated. The half effect 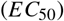 for NADPH_*ec*_ ⇒ ROS_*ec*_ (*j* = 25) is 0.838; this interaction rule upstream of IL-6 limits IL-6 to reach its maximal activity when *W*_GLU_ = 0.75. The intracellular levels of ROS were lower in GECs when stimulated with low glucose levels (below 0.5) (Fig. 13b). We observed no recovery in the intercellular gap width when exposed to all non-zero glucose levels (0.25 ≤*W*_GLU_ ≤1) (Fig. 13d). We believe other intervention strategies besides glucose control may be needed to potentially reverse the changes in gap width and loss in barrier integrity.

**Figure 13:**
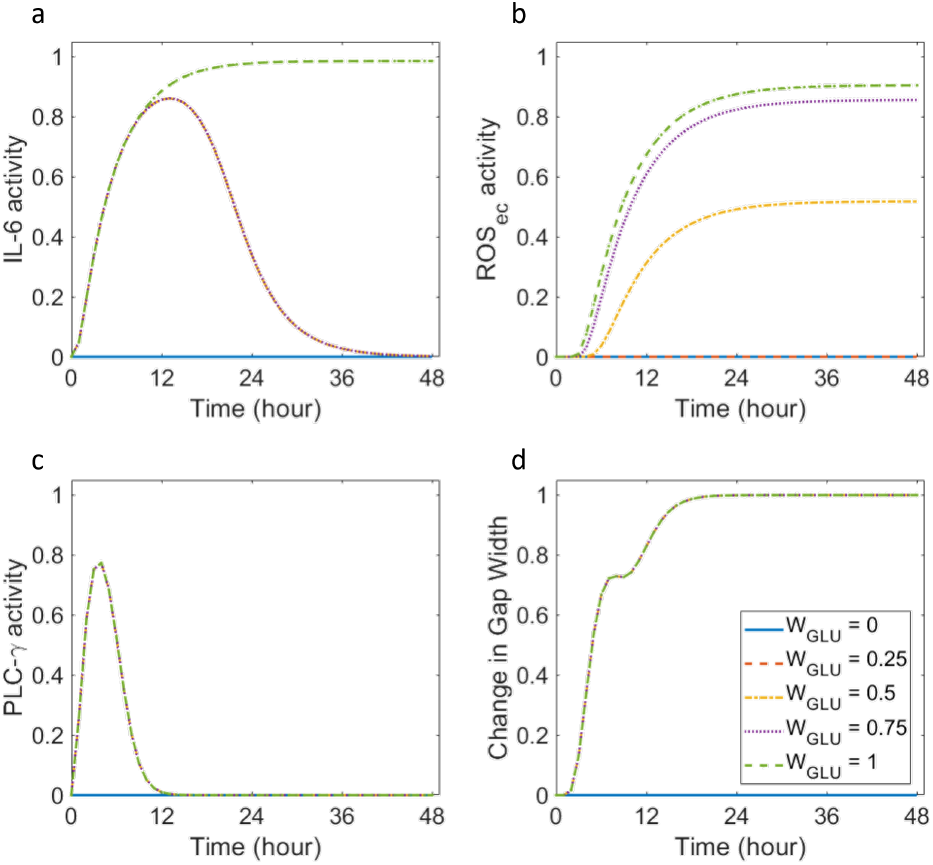
LBODEs model predictions under partially active glucose (GLU) stimulus to explore ranges of hyperglycemia in the early stage of diabetic kidney disease. *W*_GLU_ varied between 0 and 1. *W*_LPS_ was set to 0 to mimic a non-inflammatory state. All other parameters were at their nominal values for treatment with only GLU. Abbreviations are defined in Table 1.

Next, the macrophages and GECs were also stimulated with varying levels of LPS stimulus (0 ≤*W*_LPS_ ≤1) when glucose *W*_GLU_ = 1 was set as high (Fig. 14). This condition mimics the early phase of DKD development, where high glucose levels cause metabolic dysregulation in pro-inflammatory species and the environment. The nominal species parameters were based on GLU only treatment (Table 3). Increasing levels of LPS activity had minimal effect on pro-inflammatory cytokines (Fig. 14a–b) but a substantial effect on the PLC-γ upregulation (Fig. 14c). Note that 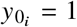 for NO for the GLU only treatment at the nominal parameter values (Fig. 4), and 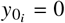 for NO for the cases of LPS only and both GLU and LPS treatments. When considering the partial activation of LPS, we selected 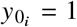 for NO to compare to the GLU only treatment dynamics (Fig. 4g) as *W*_GLU_ = 1 in all the cases shown. In contrast, different levels of partial LPS activation were considered. We observed that glucose-dependent downregulation of NO activity recovered to the baseline level as LPS activity increased above 50% (*W*_LPS_ ≥ 0.5) (Fig. 14d).

**Figure 14:**
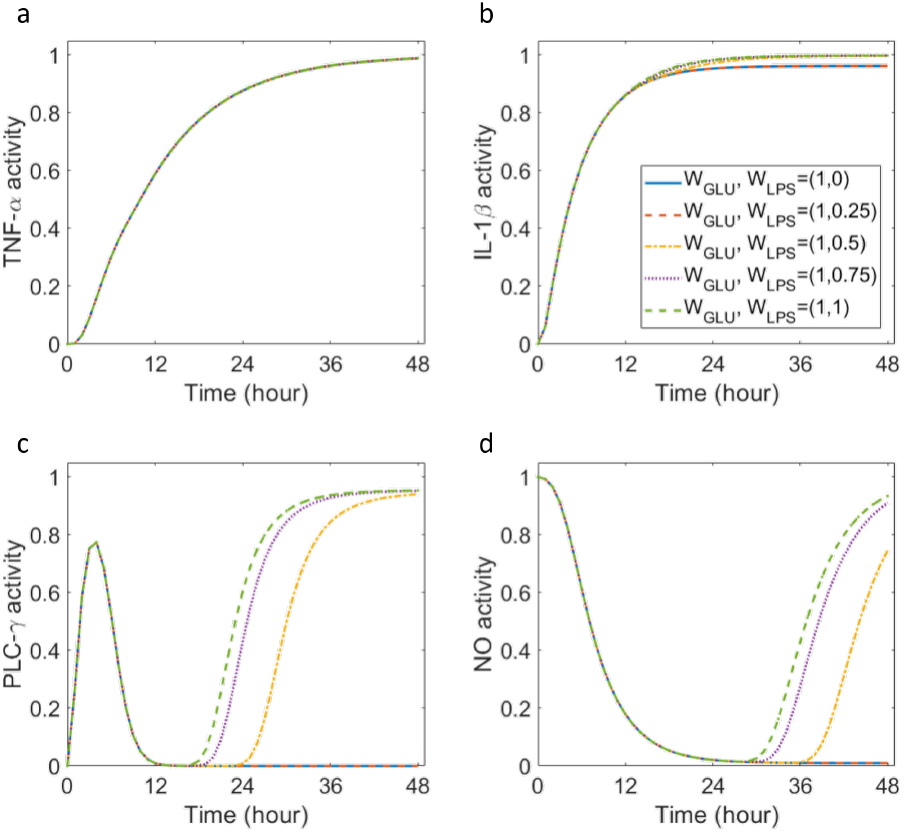
LBODEs model predictions under partially active lipopolysaccharide (LPS) and fully active glucose (GLU) stimulus (*W*_GLU_ = 1) to mimic gradual rise in inflammation in the early stage of diabetic kidney disease. *W*_LPS_ was varied between 0 and 1.All other parameters were at their nominal values for treatment with only GLU. Abbreviations are defined in Table 1.

We identified influential interactions (reaction rules) in the PSN that were sensitive to *in silico* knockdowns by reducing the reaction weight (*W*_*j*_) of all interactions and performing a local sensitivity analysis on the LBODEs model in response to these perturbations. We also repeated the process to determine the influential species by reducing the 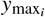 parameter of all species in the PSN. We perturbed *W*_*j*_ and 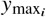 by reducing their optimal (nominal) values one-at-a-time by 15%. The optimal values for *W*_*j*_ are available in Table 2, and nominally 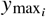 for each species *i. W* _*j*_ < 1 denotes a reduced strength of the interaction (*j*). 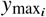 < 1 denotes the knockdown of a species (*i*) in the PSN. The sensitivity indices were calculated using *Eq. 12* for each interaction and species upon local perturbation in *W*_*j*_ and 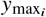,respectively. We used heatmaps to display the resulting sensitivity indices at 48 hours (Supplemental Fig. S7). We considered the nominal parameters for this analysis to be those for the treatment with both GLU and LPS (Tables 2 and 3). A magnitude of sensitivity index greater than 0.1 was used as a criterion to identify sensitive interactions and species from Supplemental Fig. S7 corresponding to a specific row labeled on the *y*-axis. In Fig. 15, we observed the predicted effects on Gap Width from targeting these sensitive interactions and species using an *in silico* knockdown test. The strengths (*W*_*j*_) of the targeted interactions and the maximal activities 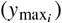 of the targeted species were reduced by 15% (Fig. 15a–b) and 50% (Fig. 15c–d) from the nominal values. For all examined interactions and species, a 15% reduction in parameter values positively correlated with the decrease in GEC Gap Width compared to the no inhibition case (no perturbations) (Fig. 15a–b). We found that 50% inhibition of the following reactions reduced the GEC Gap Width substantially (Fig. 15c): VEGF-A_mRNA_ ⇒ VEGF-A (*j* = 17), VEGF-A ⇒ VEGFR1 (*j* = 19), VEGFR1 ⇒ PLC-γ (*j* = 28), PLC-γ ⇒ Ca (*j* = 37), Ca ⇒ pJunction (*j* = 38), and pJunction Gap Width (*j* = 39). Targeting VEGFR1, NFκB, NO, PLC-γ, VEGF-A, pJunction, and Ca species in the network substantially recovered the GEC Gap Width (Fig. 15d). Among the sensitive reactions and species identified in Supplemental Fig. S7, 50% inhibition of NFκB ⇒ VEGF-A_mRNA_ reaction (*j* = 16) and VEGF-A_mRNA_ did not reduce the Gap Width or even follow the trajectory for the case of no inhibition. Instead, the predicted values of Gap Width plateau at 1 in these cases. We postulate that 50% or more inhibition of NFκB ⇒ VEGF-A_mRNA_ reaction or of VEGF-A_mRNA_ results in an imbalance in NO and Ca activity due to the negative feedback loop between NO and Ca. This imbalance can explain the increase in Gap Width slightly compared to the value without inhibition (Fig. 15c–d).

**Figure 15:**
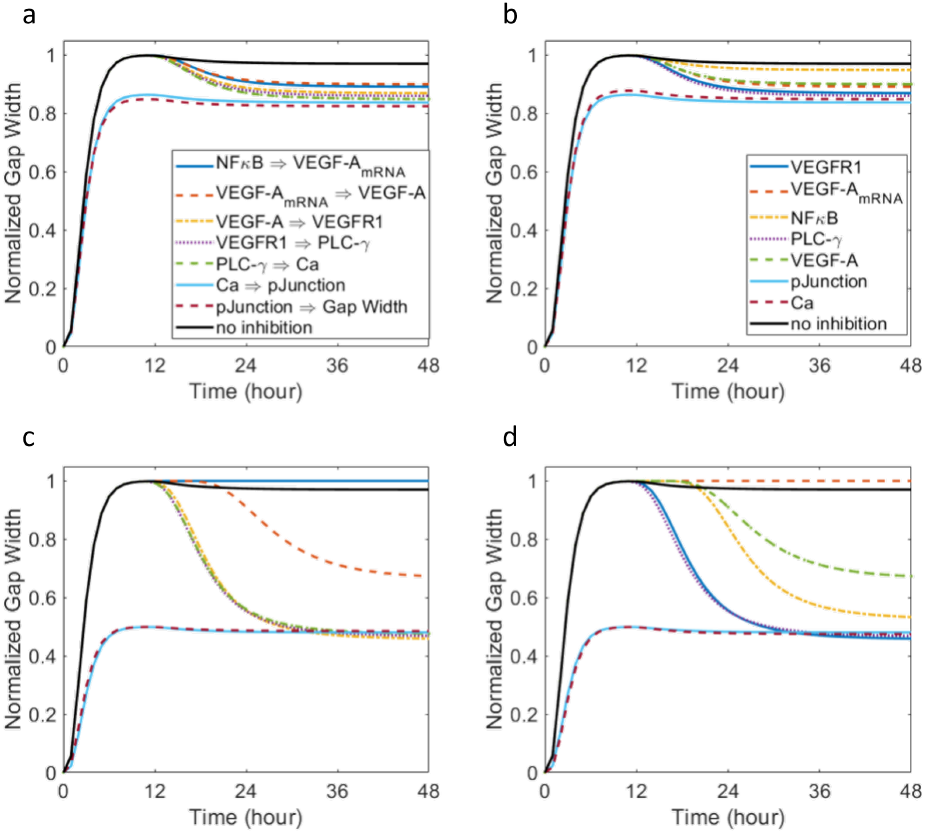
LBODEs model predictions of the normalized Gap Width between GECs under *in silico* knockdowns. Reaction weight *W*_*j*_ of each locally sensitive interaction was reduced by (a) 15% and (c) 50% from its nominal value. Maximum activity 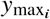 of each locally sensitive species was reduced by (b) 15% and (d) 50% from its nominal value. All other parameters were at their nominal values for treatment with both GLU and LPS. The solid black curve represents the normalized Gap Width without any changes in reaction or species parameters from their nominal values. Panels a and c share a legend, and panels b and d share a legend. Abbreviations are defined in Table 1.

## DISCUSSION

In this work, we predicted the continuous signaling dynamics of proteins in macrophages and GECs using an LBODEs modeling technique and prior knowledge of intracellular and intercellular pathways compiled into the PSN. We fitted and validated the model using published *in vitro* data. We demonstrated the effect of hyperglycemia and inflammation on signaling pathways and protein expression.

Glucose- and LPS-mediated upregulation of pro-inflammatory cytokines further augments the phenotypic switch towards the pro-inflammatory state in macrophages in the early stage of DKD (Fig. 7a–c). The model demonstrated this pro-inflammatory phenotype in macrophages with an increased expression of pro-inflammatory species, such as IL-6, TNF-α, IL-1β, and ROS. The consistent secretion of these inflammatory molecules leads to a chronic shift in macrophage phenotype (112). The combined effect of glucose and LPS affected the intensity and early activation of pro-inflammatory cytokines, oxidative stress, and growth factor activity (Figures 6 and 9). The model predicted a slight decrease in TNF-α expression after 6 hours (Fig. 8b). Multiple reactions and reaction parameters were responsible for TNF-α activity. NFκB transcription factor in macrophages and NFκB_ec_ transcription factor in GECs were responsible for direct activation of TNF-α in the network. The differences in the optimal reaction weight (*W*_*j*_) and half-maximal effect 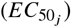 parameters (Table 2) for NFκB ⇒ TNF-α and NFκB_ec_ ⇒TNF-α can explain the shift in TNF-α activity. The model predictions demonstrated the effect of high glucose on early signs of glomerular endothelial activation through reduced VEGF-mediated kinase activity (Fig. 10d), reduced NO bioavailability (Fig. 7f) due to uncoupling of NO from its pathway, increased inflammation (Fig. 7), and loss of intercellular junction integrity (Fig. 10e). The model predicted that VEGF-A primarily regulated endothelial cell survival, kinase activity, NO homeostasis, and permeability, as seen by early PI3K_*ec*_ activation (113) (Fig. 12d), followed by leukocyte migration and inflammation mediated by VEGFR1/PLC-γ activation (52) (Fig. 12d). Besides VEGF-A, GECs can also be activated by angiopoietins, chemokine ligands, endothelin-1, and pro-inflammatory cytokines (IL-6, TNF-α, IL-1β) upon cell-cell communication with glomerular cells like podocytes (114). The proposed PSN can be extended to include other sources of GEC activation and interaction with glomerular cells. We demonstrated the effects of VEGF-A and its receptors on cell-cell junction opening. From a molecular standpoint, the endothelial barrier is sealed by cell-cell junction proteins, among which VE-cadherin serves as a cornerstone (115). The endothelial barrier is maintained by VE-cadherin and associated catenins in a non-phosphorylated state (26, 72, 116). The modulated phosphorylation of these junction proteins allows for the selective opening or closing of the intercellular barrier (11). The relationship between VEGF-dependent NO and calcium and phosphorylated junction proteins was captured as a negative feedback loop in the network. A negative feedback loop between NO and Ca allowed for better information processing and cell behavior regulation in the PSN. We observed that lower than healthy NO levels in GECs affected barrier integrity reversibly (40) (Fig. 10e). High glucose and inflammation caused a prolonged and consistent disruption of the cell-cell barrier integrity in GECs (Fig. 12e).

Regulating glucose levels and inflammation to baseline values effectively reduced cytokine expression and oxidative stress and restored NO bioavailability (Figures 13 and 14). We showed that targeting the mechanistic pathway that links VEGF-A, Ca, and phosphorylated junctions is relevant to understanding the early signs of glomerular endothelial activation and dysfunction. We identified that NFκB transcription factor in macrophages, VEGFR1, PLC-γ, Ca, and junction proteins on GECs are potential targets to control intercellular Gap Width (Fig. 15d). Further *in silico* experiments can suggest the minimum percentage of protein knockdown or reaction inhibition required to maintain intercellular GEC Gap Width. We also observed that excessive knockdown of a protein or a reaction could result in an undesirable effect on Gap Width due to the complex nature of intracellular signaling. Although we modeled the change in Gap Width using a general form of the normalized-Hill type ODE, we obtained a reasonable and qualitative prediction of change in Gap Width (Fig. 15). Our understanding of modeling the Gap Width change is limited by literature, *in vitro* data, and other mechanistic models. Further improvements to the model can be made as new information becomes available. Our predictive findings suggest that a multi-targeted approach is necessary to modulate the early markers of DKD development and progression. Multi-targeted strategies may involve glucose control (Fig. 13), attenuating inflammation (Fig. 14), reducing macrophage infiltration and transmigration via PLC-γ, and maintaining GEC barrier integrity. The potential targets identified by the model can be further analyzed and validated through experimental studies and quantitative systems pharmacology models.

Future extensions of this work could include translating the LBODEs model developed here to study protein dysregulation, GEC activation, and dysfunction in diabetic mice using *in vivo* data. This would enable the study of the effects of systemic changes in glucose concentration and long-term exposure to glucose on structural and functional changes in GECs *in vivo*. The structure of GECs consists of transcellular holes, known as fenestrations, which support the glomerular filtration barrier (117, 118). Alterations in the size and density of GEC fenestrations have been recently associated with the disruption in glomerular filtration and the progression of DKD (117). Recent literature provides more understanding of damaged fenestrated endothelium like GECs in disease conditions (117–120) and suggests that—besides VEGF, NO, and calcium—other proteins such as rho-associated protein kinases and myosin light chain kinases are involved in regulating the fenestration structure in endothelial cells (22, 121–123). We aim to extend the proposed PSN by incorporating these additional hypothesized mechanisms responsible for regulating GEC fenestration structure and permeability.

The quantitative agreement between normalized model prediction and observed data shows the usefulness of the LBODEs framework in understanding complex disease mechanisms. Furthermore, the LBODEs framework provides testable hypotheses on the most sensitive mechanisms that lead to GEC activation and dysfunction in DKD. As the modeling framework is composed of ordinary differential equations, it is suitable to be used with many commercially available tools for analysis of dynamical systems and easy to integrate with other mechanistic models in systems biology (39, 124, 125). Throughout this study, we have depicted how these valuable and practical methods for identifiability, parameter sensitivity, and parameter uncertainty analysis can be integrated with the LBODEs dynamical system.

We also acknowledge some limitations of the LBODEs framework compared to other biochemical models. The species behavior is quantified as a fractional activation or inhibition rather than absolute quantities such as concentration. The LBODEs framework is limited to analyses with normalized protein levels. The presented LBODEs framework represents all protein-protein interactions using activating or inhibiting Hill functions, but other mathematical functions can be used as appropriate. The model predictions are limited to the structure of the proposed PSN, but additional processes, proteins, and pathways can be added. Despite careful consideration when pooling data from multiple experimental studies, there were some discrepancies across cell culture studies, especially when measuring oxidative stress and nitric oxide levels. ROS species are often short-lived and highly reactive, making them challenging to measure. Usually, special assays are needed to detect specific forms of oxidative stress and nitric oxide. Such discrepancies could explain the differences in observed normalized data across studies, and they limit model validation (Fig. 7e). Furthermore, in the absence of GEC-specific experimental studies, we obtained protein concentrations from similar endothelial cell types. We believe the observed uncertainty in prediction can be resolved as more comprehensive dynamic and cell-specific data becomes available.

Despite such limitations, the present model motivates new experiments and testable hypotheses that can better elucidate the early signs of DKD progression. The LBODEs modeling framework enables quantifying protein behavior for large-scale and complex signaling networks, especially when experimental validation is limited. The mechanisms identified from the LBODEs model can be used to develop a less complex mechanistic model for which kinetic and cell-specific parameters are available. Our proposed *in silico* exploration of signal transduction and protein expression could support pre-clinical research to test new mechanisms for therapeutic intervention or improve on existing strategies.

## CONCLUSIONS

In this study, we presented a multi-cellular signaling LBODEs model to understand the role of diabetes and immune response in the progression of microvascular complications in the diabetic kidney. We created a signaling network of crosstalk between macrophages and GECs stimulated by two inputs—high GLU and a pro-inflammatory stimulus LPS. The signaling network was modeled using continuous ODEs and logic-based interaction rules, where interaction between species was formulated using normalized Hill functions. The model was fitted and validated against published data from *in vitro* experiments performed on murine macrophage cell lines and endothelial cell lines. Our modeling approach combined the advantages of quantitative biochemical models and qualitative LBMs. These pathways and interactions between macrophages and GECs in the early stage of DKD have not been mathematically studied before. In the presence of high GLU and a pro-inflammatory LPS stimulus, we observed upregulation of pro-inflammatory cytokines (IL-6, TNF-α, and IL-1β), oxidative stress, VEGF-A, and intercellular GEC gap width. We demonstrated the role of oxidative stress in macrophages in regulating NFκB transcription factor and NO bioavailability in the downstream signaling response. We found that partial knockdown of VEGFR1, PLC-γ, pJunction, and Ca have the potential to partially recover the intercellular GEC barrier integrity. The potential protein and interaction targets can be further tested through modeling or experimentation. Moreover, the proposed PSN and model can be translated to study histological changes in GECs *in vivo*. Through our signaling network and LBODEs model, we have studied glucose-mediated inflammatory pathways in the diabetic kidney to support an understanding of DKD development and progression.

## APPENDIX

### Model equations

The following LBODEs (*Eqs. A1 –A30*) govern the PSN interactions. The species abbreviations are defined in Table 1. The parameter values are available in Tables 2 and 3. Other model term definitions are available in *Eqs. 1 – 7*.

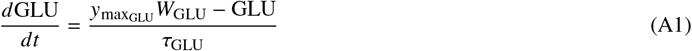

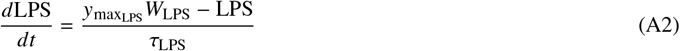

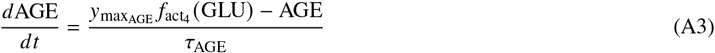

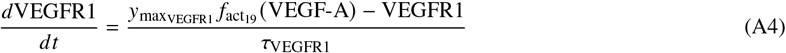

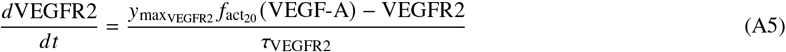

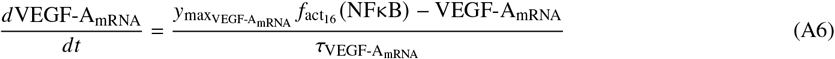

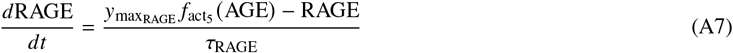

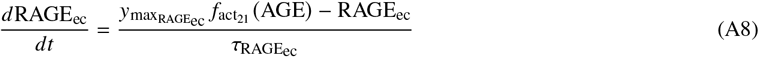

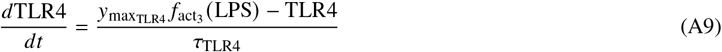

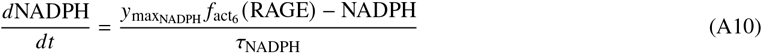

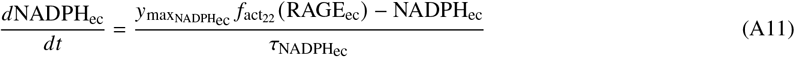

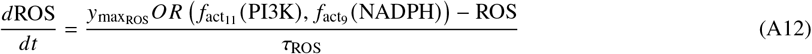

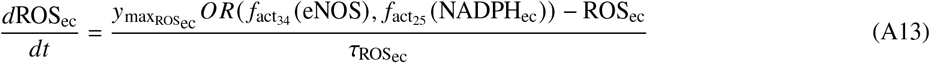

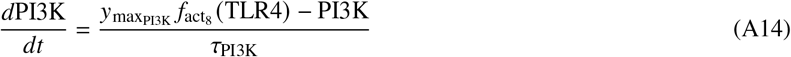

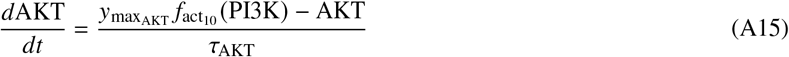

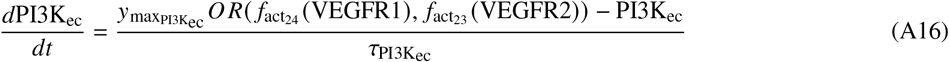

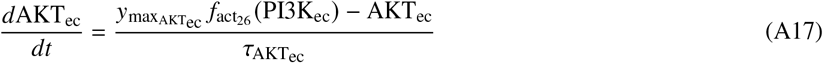

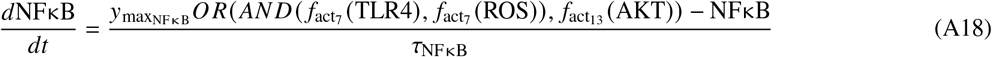

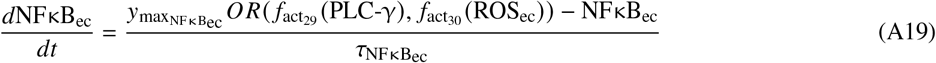

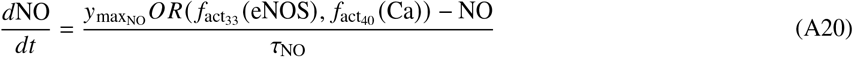

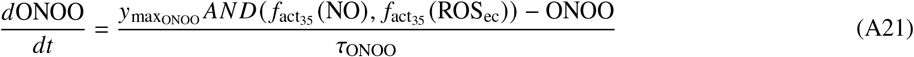

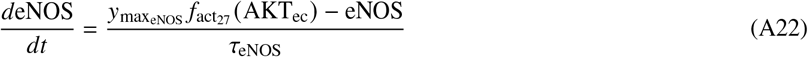

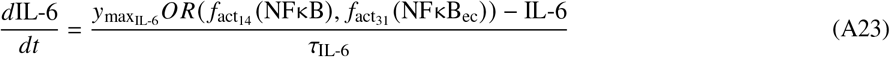

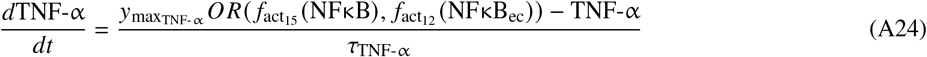

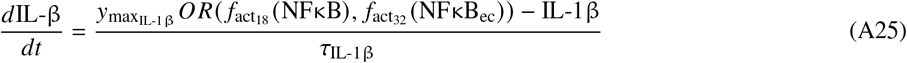

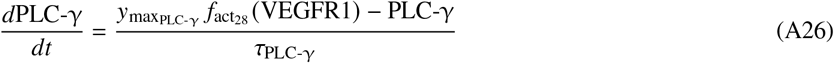

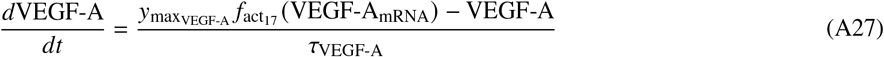

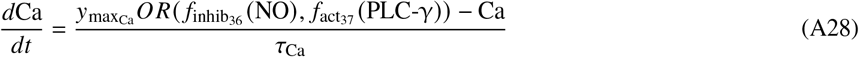

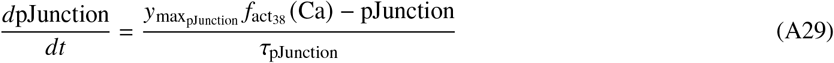

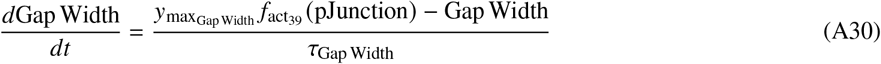

## SUPPLEMENTAL MATERIAL

Supplemental Figs. S1–S7.

## DATA AVAILABILITY

No primary data were generated in this study. The supporting MATLAB code, including parameter values, scripts for plotting, data, and documentation, can be accessed through an open-source GitHub repository (126) available at this link: https://github.com/ashleefv/KidneyImmuneLBM

## ACKNOWLEDGMENTS

We thank lab members for their thorough feedback on this manuscript. Preprint (127) is available at https://doi.org/10.1101/2023.04.04.535594.

## AUTHOR CONTRIBUTIONS

ANFV and KP conceived and designed research, KP performed experiments, KP analyzed data, ANFV and KP interpreted results of experiments, ANFV and KP prepared figures, KP drafted the manuscript, ANFV and KP edited and revised manuscript, and ANFV and KP approved the final version of the manuscript.

