## Supplemental Material for "Logic-Based Modeling of Inflammatory Macrophage Crosstalk with Glomerular Endothelial Cells in Diabetic Kidney Disease"

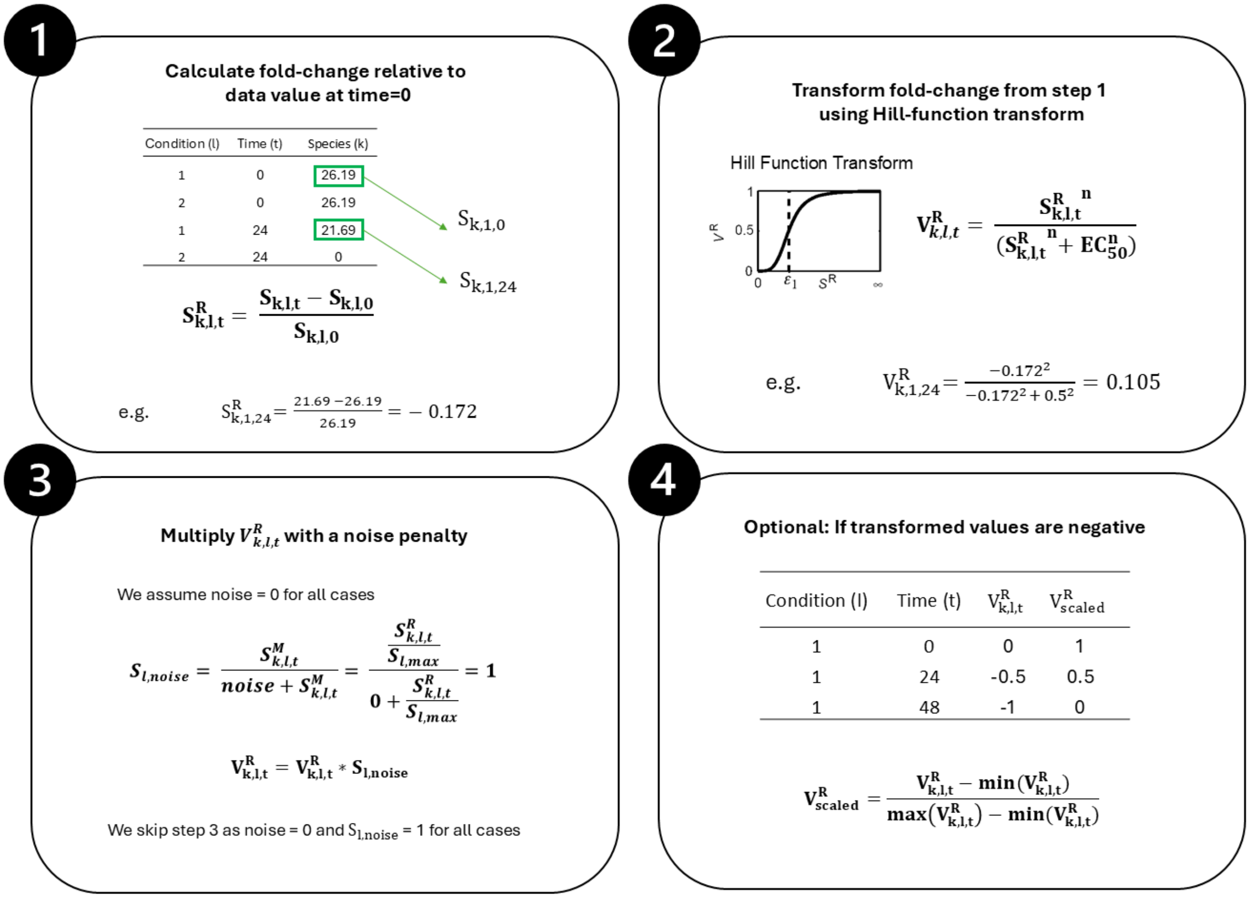


Figure S1: Steps involved in the data normalization method in CellNOptR. Step 1. Data values for a given species (k) are transformed to a fold change for the same experimental treatment condition (l) at time (t) = 0. Step 2. The fold change is transformed using a Hill function. Step 3. A penalty for noise in the data (if any) is multiplied by the transformed value to compute the normalized data value. Step 4. If Hill-transformed fold changes after noise penalty have negative values, the normalized data are rescaled using the minimum and maximum values in the normalized data list. $S_{k,l,t}$: data value for species (k) at time (t) and condition (k). $S^R_{k,l,t}$: fold-change for a species (k) at time (t) and condition (l). $V^R_{k,l,t}$: Hill-transformed value. $EC_{50} = 0.5$. $n = 2$. $S_{l,noise}$: noise penalty. $S_{l,max}$: maximum of the values for a condition (l). $V^R_{scaled}$: negative Hill-transformed values scaled to positive values.

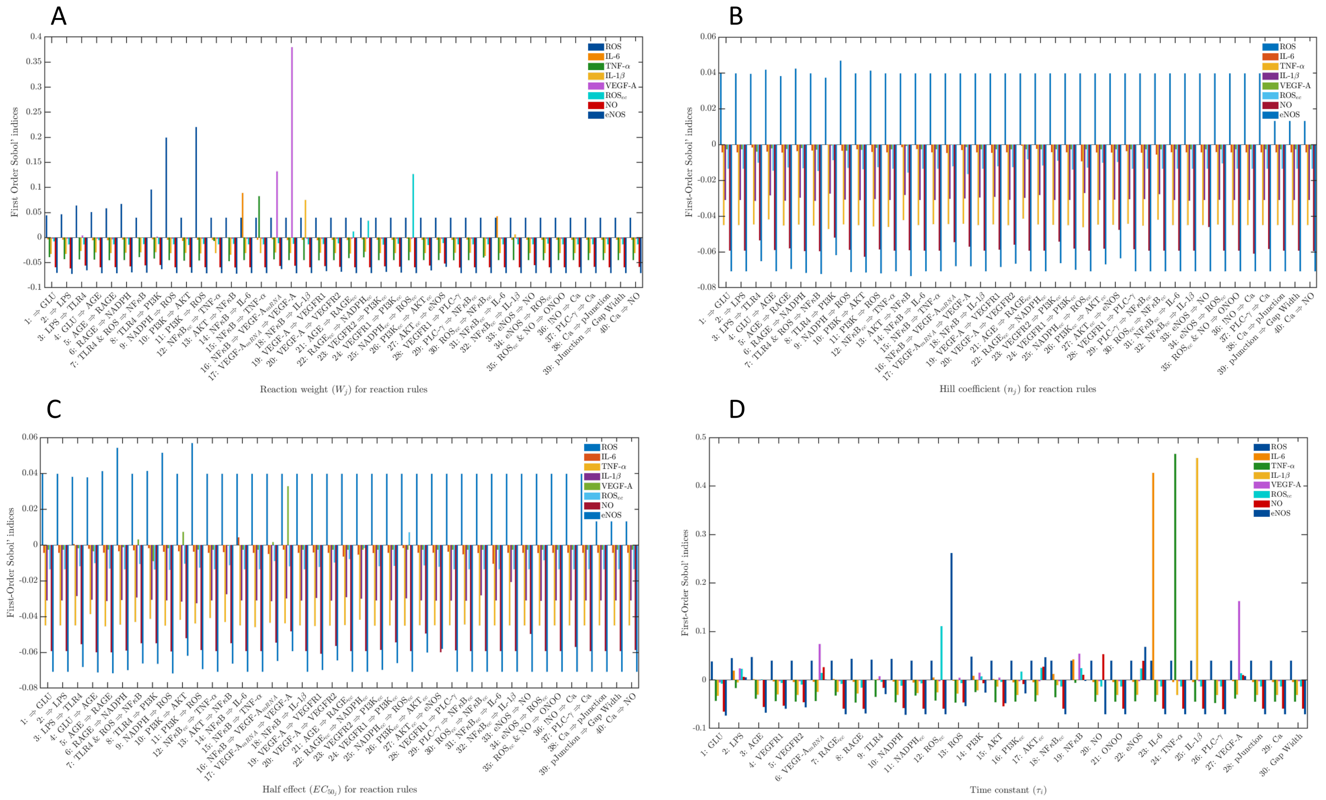


Figure S2: First order Sobol indices for (A) reaction weight ($W_j$), (B) Hill coefficient ($n_j$), (C) half effect ($EC_{50_j}$) for respective reaction rules, and (D) time constant ($\tau_i$) for respective species were calculated using the UQLab toolbox. The Sobol sensitivity for each parameter was calculated against species response of ROS, IL-6, TNF-α, IL-1β, VEGF-A, $ROS_{ec}$, NO, and eNOS at 48 hours. These species are ordered on the plot from left to right for each parameter. Abbreviations are defined in Table 1 in the main text.

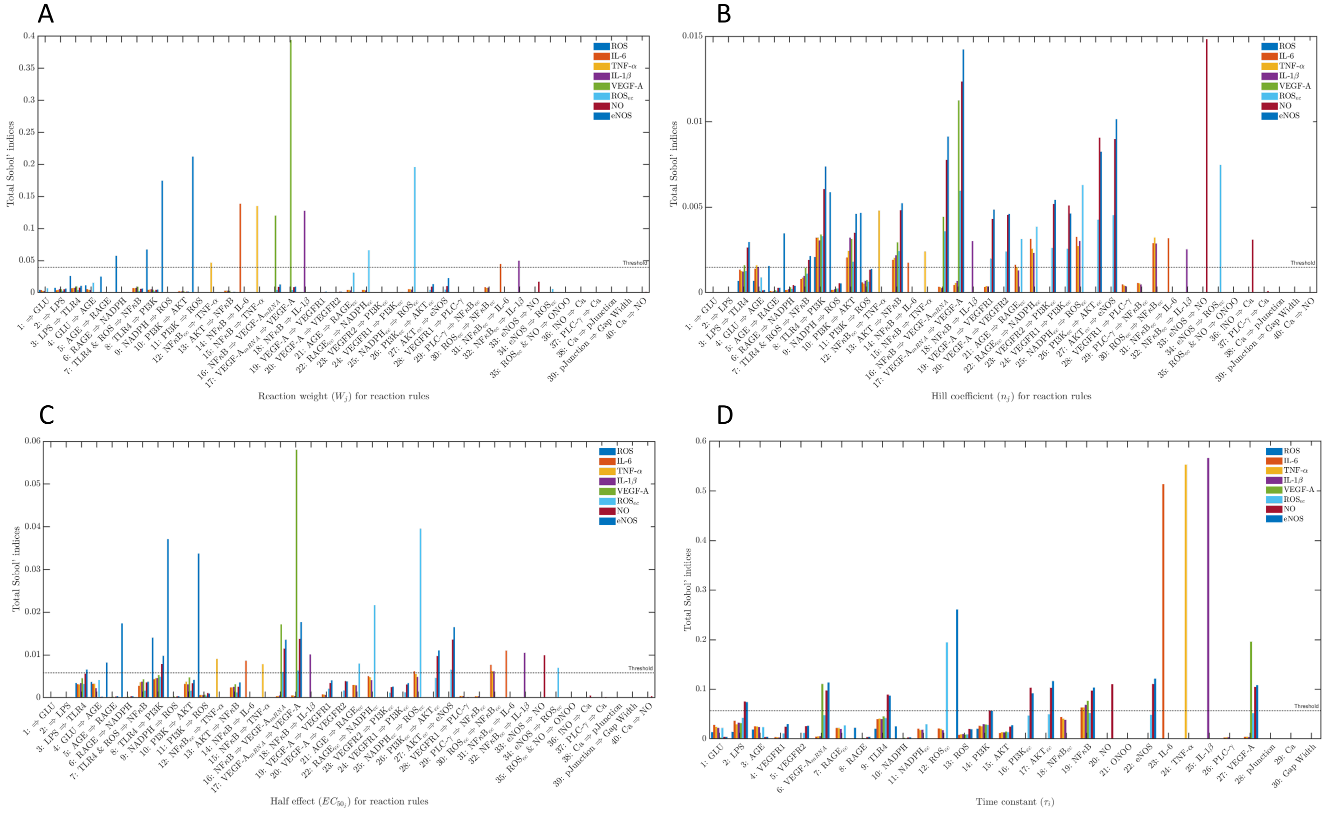

Figure S3: Total order Sobol indices for reaction weight ($W_j$), (B) Hill coefficient ($n_j$), (C) half effect ($EC_{50_j}$) for respective reaction rules, and (D) time constant ($\tau_i$) for respective species were calculated using the UQLab toolbox. The Sobol sensitivity for each parameter was calculated against species response of ROS, IL-6, TNF-α, IL-1β, VEGF-A, $ROS_{ec}$, NO, and eNOS at 48 hours. These species are ordered on the plot from left to right for each parameter. Abbreviations are defined in Table 1 in the main text.

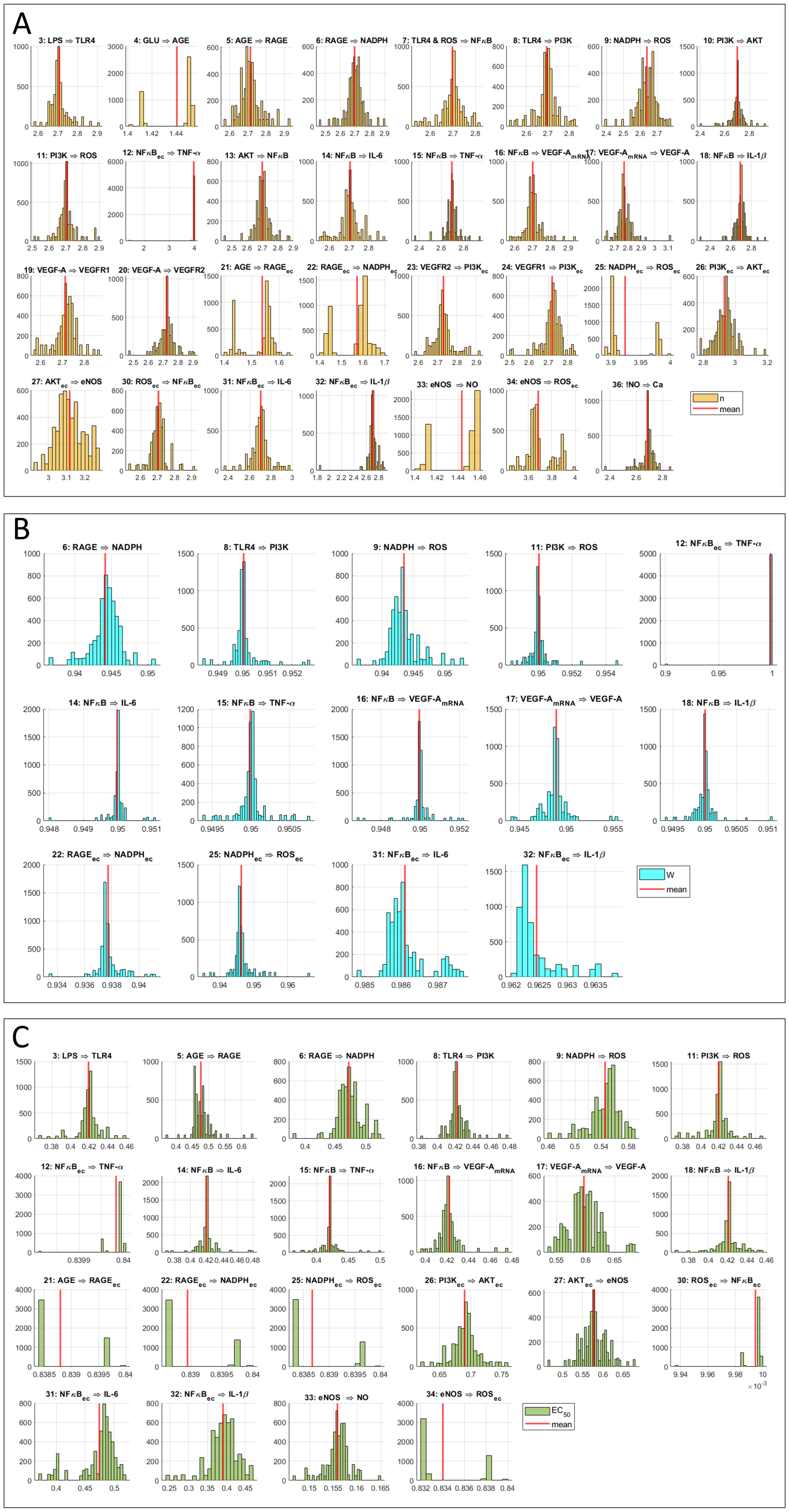


Figure S4: The posterior distribution of acceptable reaction parameter values. The histograms represent the (A) reaction weight ($W_j$) (blue), (B) Hill coefficient ($n_j$) (yellow), and (C) half-maximal effect ($EC_{50_j}$) (green) for the respective reaction index $j$ and rule as reported in Table 2. The red vertical line represents the nominal parameter values used for model prediction. The reaction index $j$ and the reaction rule are provided as the title for each subfigure. Abbreviations are defined in Table 1 in the main text.

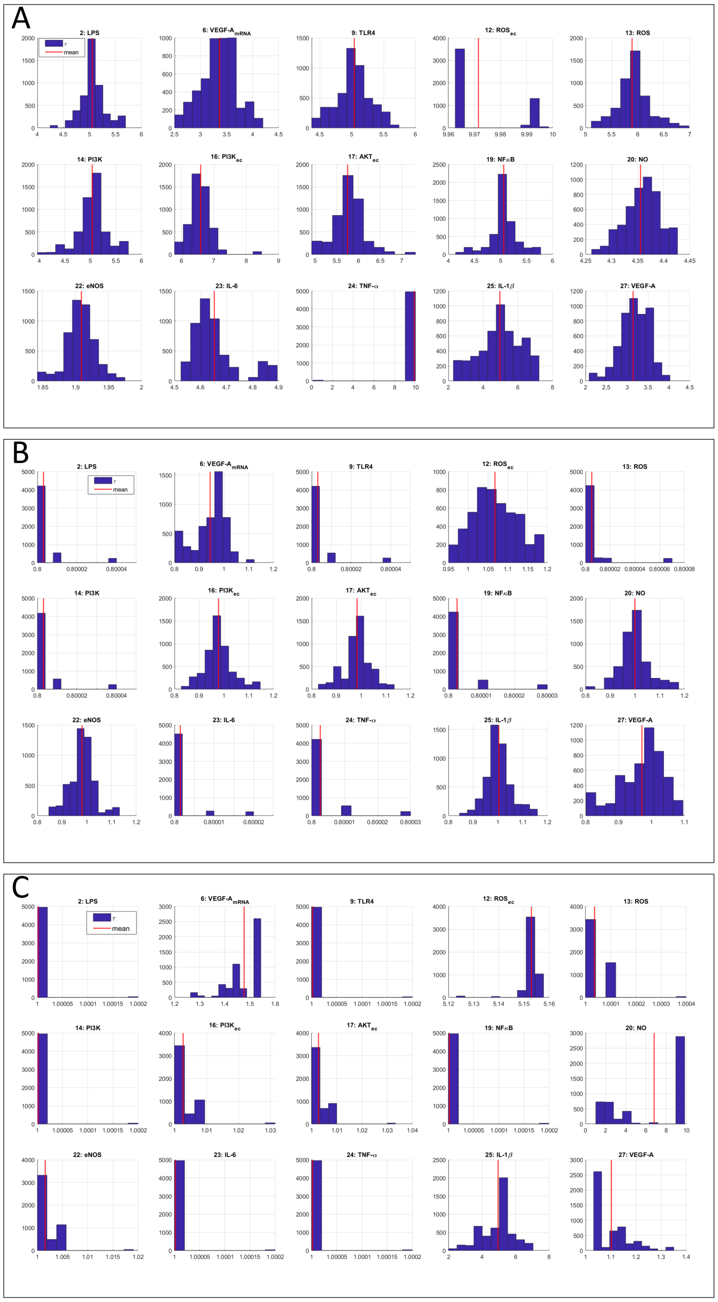


Figure S5: The posterior distribution of acceptable species time constant values for the treatments: (A) GLU only, (B) LPS only, and (C) both GLU and LPS. The histograms (navy blue) represent the time constant ($\tau_i$) parameter for the respective species index $i$ as reported in Table 3. The red vertical line represents the nominal parameter values used for model prediction. The species index $i$ and species name are provided as the title for each subfigure. Abbreviations are defined in Table 1 in the main text.

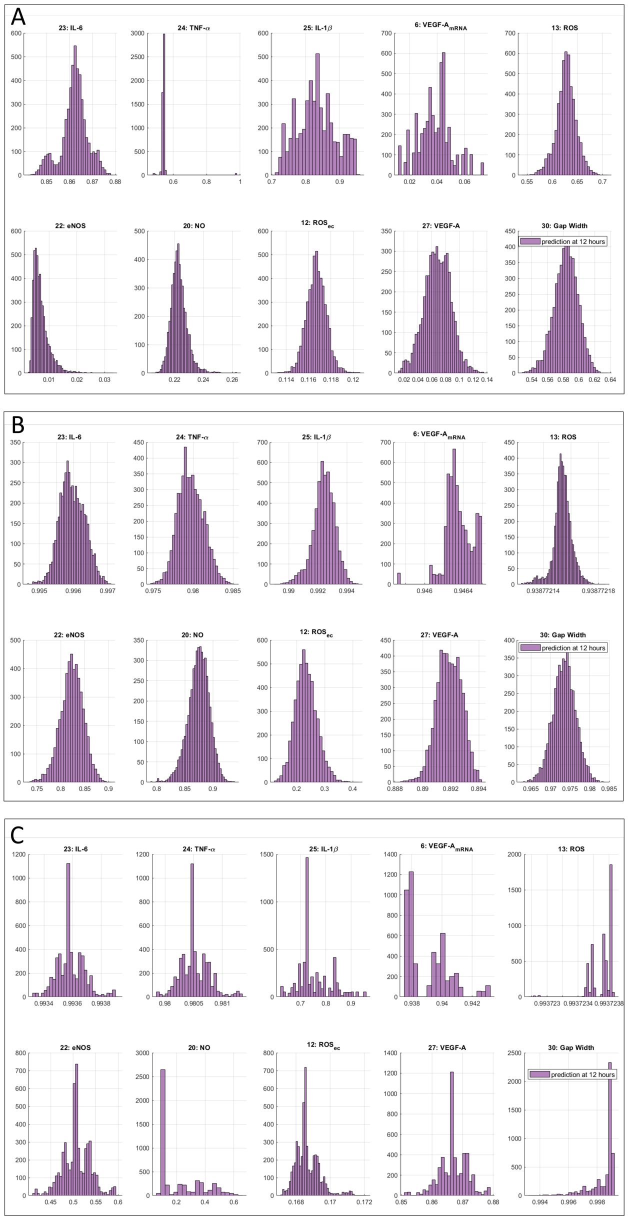


Figure S6: The posterior distribution of prediction responses at 12 hours for the IL-6, TNF-α, IL-1β, VEGF-A$_{mRNA}$, ROS, eNOS, NO, ROS$_{ec}$, VEGF-A, and Gap Width for the treatments: (A) GLU only, (B) LPS only, and (C) both GLU and LPS. The species index $i$ and species name are provided as the title for each subfigure. Abbreviations are defined in Table 1 in the main text.

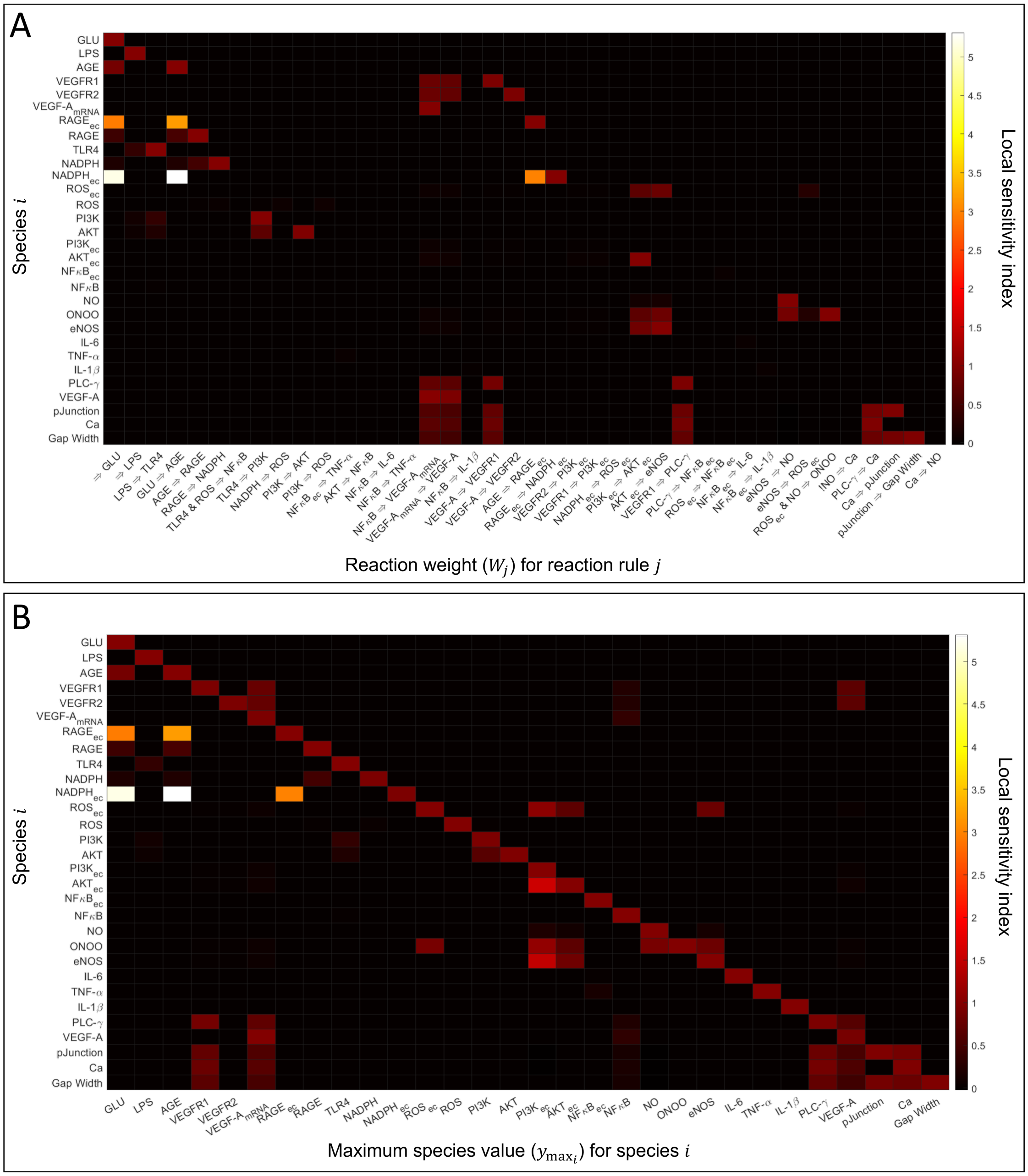


Figure S7: *In silico* perturbations in reaction weight and maximal activity using local sensitivity analysis with 15% decreases of parameters one at a time from their optimal values. The local sensitivity index of change in output species $i$ (vertical axis) against the change in (A) $W_j$ parameters of respective reactions (horizontal axis) and (B) $y_{\max_i}$ parameters of respective species (horizontal axis). The local sensitivity index was calculated using *Eq. 12*. All other parameters were at their nominal values for treatment with both GLU and LPS. The color bar indicates the magnitude of the sensitivity index. The subscript "ec" denotes intracellular expression within glomerular endothelial cells in the network. Abbreviations are defined in Table 1 in the main text.
